# Enhancement of hippocampal-thalamocortical temporal coordination during slow-frequency long-duration anterior thalamic spindles

**DOI:** 10.1101/2021.10.03.462943

**Authors:** Zahra Alizadeh, Amin Azimi, Maryam Ghorbani

## Abstract

Temporal nesting of cortical slow oscillations (SO), thalamic spindles and hippocampal ripples indicates the succession of regional neuronal interactions required for memory consolidation. However how the thalamic activity during spindles organizes hippocampal dynamics remains largely undetermined. We analyzed simultaneous recordings of anterodorsal thalamus and CA1 in mice to determine the contribution of thalamic spindles in cross-regional synchronization. Our results indicated that temporal hippocampo-thalamocortical coupling were more enhanced during slower and longer thalamic spindles. Additionally, spindles occurring closer to SO trough were more strongly coupled to ripples. We found that the temporal association between CA1 spiking/ripples and thalamic spindles was stronger following spatial exploration compared to baseline sleep. We further developed a hippocampal-thalamocortical model to explain the mechanism underlying the duration and frequency-dependent coupling of thalamic spindles to hippocampal activity. Our findings shed light on our understanding of the functional role of thalamic activity during spindles on multi-regional information transfer.

## Introduction

The precise temporal relationship of cortical slow oscillations (SOs), thalamic spindles and hippocampal ripples which are three main NREM rhythms are believed to be important for memory consolidation during sleep (Helfrich et al., 2018; Latchoumane et al., 2017; Maingret et al., 2016; Niknazar et al., 2015). Especially the mediating role of thalamic induced spindles in the hippocampal-neocortical dialogue essential for memory processing has been proposed by previous studies (Diekelmann and Born, 2010; Fogel and Smith, 2011; Latchoumane et al., 2017; Ngo et al., 2020). However how spindles with differential properties contribute to the multi-regional communication still remains poorly understood. Although the spindles are generated in the thalamus and transferred to cortex via the thalamocortical projections (Timofeev and Bazhenov, 2005), the existing literature providing evidence for spindle-ripple coupling focused on the spindle activity detected from the cortex or hippocampus (Clemens et al., 2011; Helfrich et al., 2019; Jiang et al., 2019; Mölle et al., 2006; Ngo et al., 2020; Siapas and Wilson, 1998; Sirota et al., 2003; Staresina et al., 2015; Varela and Wilson, 2020). Recent studies revealed modulation of the thalamic units to the hippocampal ripples (Yang et al., 2019; Viejo and Peyrache, 2020; Varela and Wilson, 2020), but the precise temporal association of CA1 units and thalamic spindles is still unknown. A recent study reported stronger spike-filed coupling during spindles in anterodorsal (AD) thalamus compared to barrel cortex (Bandarabadi et al., 2020). In addition, spindles detected from the thalamic LFP were more strongly coupled to the slow waves than the ones detected from the cortical LFP. Due to strong anatomical projections from the thalamus to the hippocampus, it is possible that thalamic spindles also drive the hippocampal activity directly. Hence given the latency between the thalamic and cortical LFPs during the spindles (Mak-McCully et al., 2017) and their differential coupling to the unit activity (Bandarabadi et al., 2020), it is important to investigate the precise temporal coordination of hippocampal unit activity/ripples and the spindles detected from the thalamus.

Here, we addressed this gap by characterizing the fine temporal coordination between thalamic spindles and hippocampal activity using simultaneous recordings of AD thalamus and CA1 (single units and LFPs) in freely moving mice before and after spatial exploration. We specifically tested the hypothesis that the hippocampal-thalamocortical temporal coordination depends on the oscillating frequency and duration of the thalamic spindles as well as their locking phase to SOs. A wide frequency range of 7-15 Hz are typically considered for spindles in rodents. Within this frequency band in human sleep slow and fast spindles with differential properties and functional relevance for memory consolidation are discriminated (Fernandez and Lüthi, 2020; Klinzing et al., 2016; Ayoub et al., 2013; Dehnavi et al., 2021; Hashemi et al., 2019). We hypothesize that the precise temporal modulation of CA1 units/ripples with spindles is frequency dependent so that spindles with lower frequencies provide longer windows of opportunity for cross-regional synchronization and information transfer. We further developed a simplified hippocampal-thalamocortical neural mass model that can spontaneously generate SOs, slow/fast spindles and hippocampal ripples to investigate the mechanisms underlying multi-regional temporal interactions. Our findings provide evidence for the functional role of slow long-duration thalamic spindles in the hippocampal-thalamocortical interaction required for successful memory consolidation.

## Results

### AD neurons were more strongly coupled to slow than fast spindles

We used simultaneous recordings of AD thalamus and CA1 (single units and LFPs) in freely moving mice to investigate the mediating role of thalamic spindles in multi-regional interactions during SWS. Thalamic and CA1 LFFs were used to detect spindles/SOs and ripples (150-200 Hz) respectively (Figure 1A). We hypothesized that spindles with lower frequencies provide a longer temporal window for hippocampal synchronized activity enhancing multi-regional communication. Hence in this study, slow (7-10 Hz) and fast (11-15 Hz) spindles were detected and analyzed separately (Figure 1B). To explore the differential properties of slow versus fast spindles and avoid the overlapping fast and slow spindles (Mölle et al., 2011; Andrillon et al., 2011; Gardner et al., 2013), we considered only isolated fast and slow spindles excluding the fast (slow) spindles that were accompanied by slow (fast) spindles (search window of ± 0.5 s around the spindle onset and offset). There was no considerable difference between the density, duration or amplitude of AD thalamic slow and fast spindles (mean ± SD, slow spindle: density = 0.22 ± 0.52 s^−1^, duration = 1.06 ± 0.62 s, amplitude = 3.62 ± 0.93 SD, n =190006, fast spindle: density = 0.21 ± 0.51 s^−1^, duration = 1.05 ± 0.61 s, amplitude = 3.71 ± 0.92 SD, n = 180175). To investigate the modulation of AD neurons with thalamic spindles, we computed the z-score normalized AD unit activity during fast and slow spindles (Figure 1C-1D; see Materials and methods for details). AD unit activity increased around the peak of both slow and fast spindles which is consistent with a recent study (Bandarabadi et al., 2020). We next calculated the modulation index as the difference between the peak to trough of the curve of each peri-spindle-peak spike histograms ± 0.1 s around time 0. This index was significantly larger for slow versus fast spindles indicating of a stronger modulations of AD neurons to slow than fast spindles (Figure 1E, p = 4.93 × 10^−52^, Wilcoxon signed rank test, mean ± SD, slow spindle: MI = 5.4 ± 0.05 z-score, fast spindle: MI = 5.1 ± 0.05 z-score, n = 464). We further studied the phase-locking of AD units to thalamic spindles by computing the phase of Hilbert transform of filtered thalamic LFP in the spindle frequency range at the time of the spikes of AD units occurring ± 0.5 s around the spindle peaks. 52.3 % and 42.2% of the AD units were phase-locked to slow and fast spindles respectively (p < 0.05, Rayleigh test, n = 464). Across all units, the preferred phases were also non-uniformly distributed for the slow and fast spindles so that for most of the cells the preferred phase of spike occurrence was between −90° and +90°, i.e., around the spindle peak, 0° (Figure 1F, p = 3.87 × 10^−33^, z = 71.65, Rayleigh test, mean ± SD, slow spindle: phase = − 21.73° ± 38.76°, fast spindle: p = 2.16 × 10^−32^, z = 70.04, Rayleigh test, phase = − 57.53° ± 58.23°, n = 464). The preferred phase was also closer to the spindle peak for slow versus fast spindles (p = 2.60 × 10^−4^, circular-mtest). We next compared the strength of AD unit coupling to slow versus fast spindles by computing the resultant vector length of the phases for each unit separately for slow and fast spindles. The resultant vector length was larger for slow spindles compared to the fast spindles (Figure 1G, p = 0.001, Wilcoxon signed rank test, mean ± SD, slow spindle: resultant vector length = 0.06 ± 0.05, fast spindle: resultant vector length = 0.05 ± 0.05, n = 464) confirming stronger phase-locking of AD units to slow than fast spindles.

**Figure 1.**
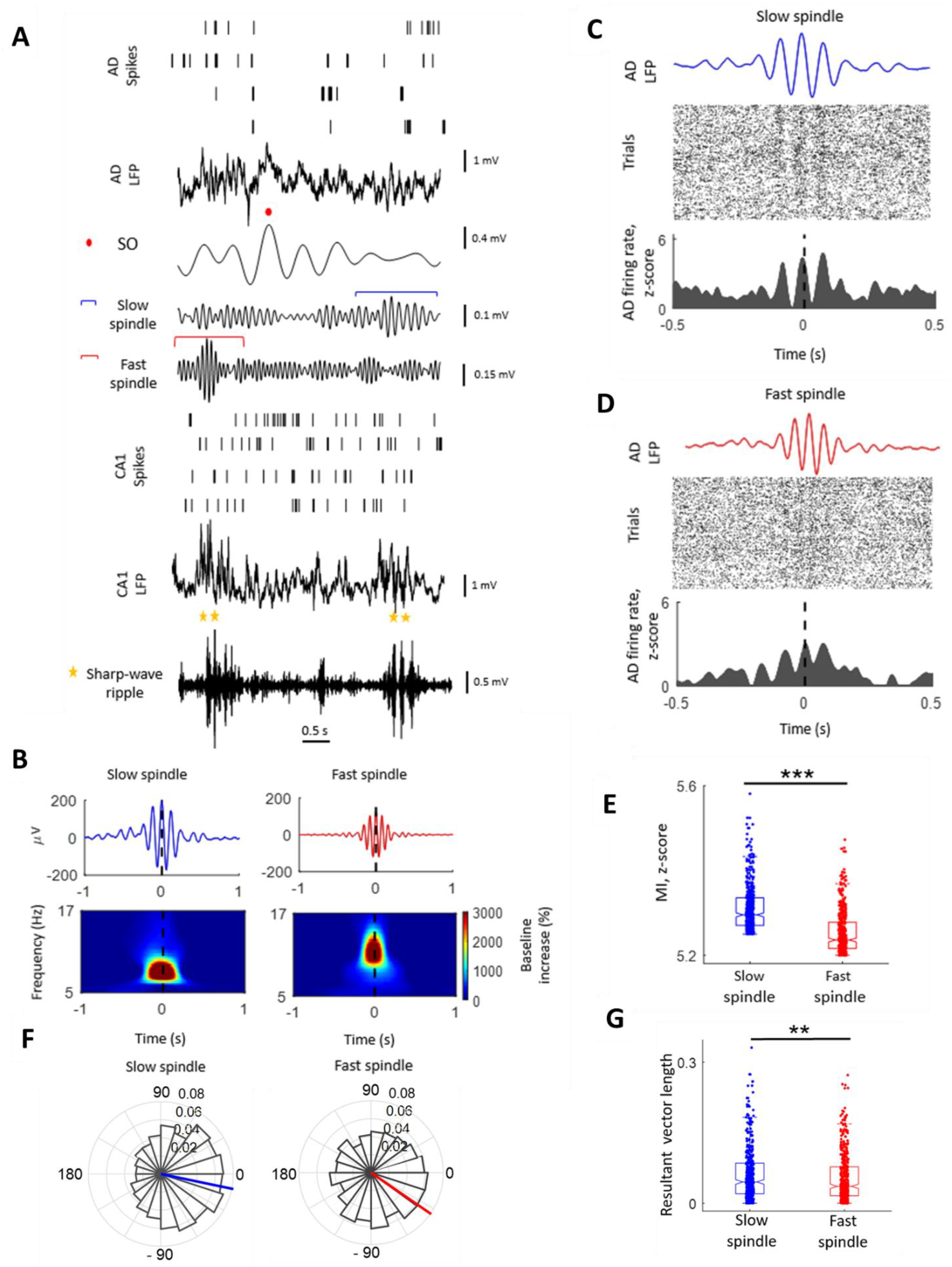
Modulation of AD neurons to thalamic fast and slow spindles. (A) Sample unit spikes and LFP traces simultaneously recorded in AD and CA1. AD LFP were used to detected SO (marked above the 0.5-2 Hz filtered AD LFP by a red circle), slow spindle (marked above the 7-10 Hz filtered AD LFP by blue bracket) and fast spindle (marked above the 11-15 Hz filtered AD LFP by red bracket). Sharp-wave ripple (marked above the 150-200 Hz filtered CA1 LFP by yellow stars) were detected using CA1 LFP. (B) Top, grand average (± SEM) of slow (Left) and fast (Right) spindles time-locked to fast and slow spindle peaks respectively. Bottom, average of slow (Left) and fast (Right) spindle peak–locked TFR (percentage change from pre-event baseline, n = 190061 and n = 180175 for slow and fast spindles respectively). (C) Top, representative average AD LFP traces filtered in the slow spindle range (7-10 Hz) aligned to the spindle peaks. Spike rasters (Middle) and peri-slow spindle-peak histograms (Bottom) of AD spikes from one unit (z-score normalized). (D) The same as (C) for fast spindles (11-15 Hz). (E) Comparison of the modulation index of AD units defined as the difference between peak to trough of the curve of each peri-spindle-peak spike histograms ± 0.1 s around time 0 for slow (blue) and fast (red) spindles. (F) Circular histograms of slow (Left) and fast (Right) spindle phases at the time of AD unit spikes, showing a preferential occurrence of spikes near spindle peak (zero degree). (G) Comparison between the resultant vector length of the phases of slow (blue) and fast (red) spindles at the time of AD unit spikes. Two asterisk indicates p<0.01, three p<0.001, Wilcoxon signed rank test, n = 464.

### Temporal correlation of CA1 units with AD spindles

We next examined the modulation of CA1 units with AD thalamic spindles by computing peri-spindle-peak spike histograms of CA1 units. Figure 2A-2B displays the strong firing modulations of CA1 units with thalamic spindles for one representative unit. The modulation index was significantly larger for slow spindles compared to the fast spindle (Figure 2C, p= 3.69 × 10^−11^, Wilcoxon signed rank test, mean ± SD, slow spindle: MI = 4.05 ± 0.05 z-score, fast spindle: MI = 3.49 ± 0.06 z-score, n = 58). To further study the temporal relationship of CA1 units with thalamic spindles, we computed the phase-locking of CA1 unit spikes to the spindles using Hilbert transform. 48.2% and 46.5% of the CA1 units were phase-locked to slow and fast spindles respectively (p < 0.05, Rayleigh test, n = 58). Across all units, the preferred phases were also non-uniformly distributed for the slow and fast spindles showing a preferential occurrence of CA1 unit spikes around the spindle peak (Figure 2D, slow spindle: p = 0.01, z = 4.26, Rayleigh test, mean ± SD, phase = 8.01° ± 35.21°, fast spindle: p = 0.02, z = 3.81, Rayleigh test, phase = − 18.2° ± 56.09°, n = 58). In addition, the resultant vector length of the phases was larger for slow spindles compared to the fast spindles (Figure 2E, p = 0.012, Wilcoxon signed rank test, mean ± SD, slow spindle: resultant vector length = 0.06 ± 0.05, fast spindle: resultant vector length = 0.04 ± 0.04, n =58). These results indicate stronger phase-locking of CA1 units to slow than fast spindles.

**Figure 2.**
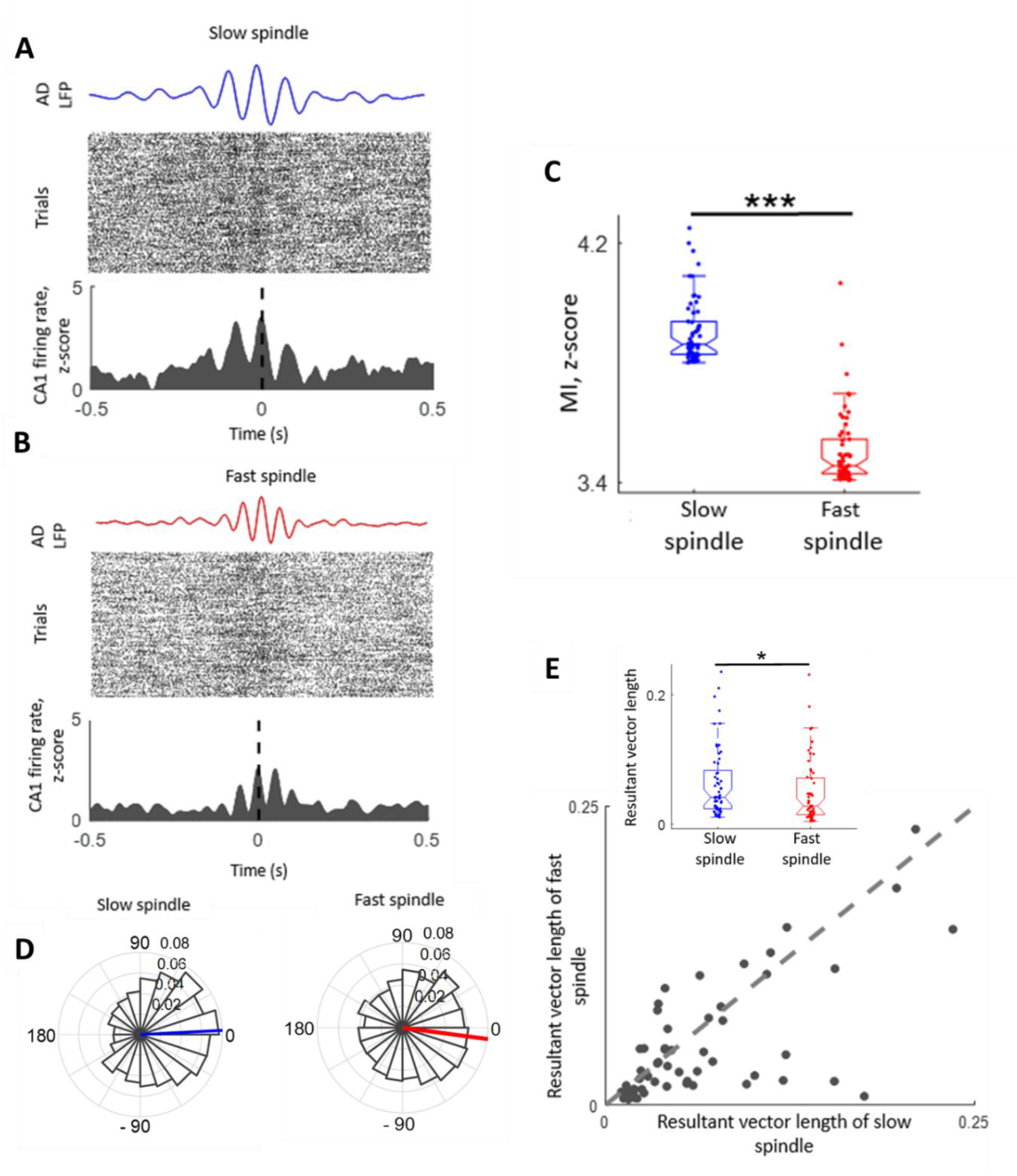
Modulation of CA1 units to thalamic spindles. (A) Top, representative average AD LFP traces filtered in the slow spindle range (7-10 Hz) aligned to the spindle peaks. Spike rasters (Middle) and peri-slow spindle-peak histograms (Bottom) of spikes from one CA1 unit (z-score normalized). (B) The same as (A) for fast spindles (11-15 Hz). (C) Comparison of the modulation index of CA1 units defined as the difference between peak to trough of the curve of each peri spindle-peak spike histograms ± 0.1 s around time 0 for slow (blue) and fast (red) spindles. (D) Circular histograms of slow (Left) and fast (Right) spindle phases at the time of CA1 unit spikes, showing a preferential occurrence of CA1 spikes near spindle peak (zero degree). (E) The resultant vector length of the fast spindle phases at the time of CA1 units versus the one for slow spindles. Each point shows one CA1 unit. Inset shows the average resultant vector length of CA1 units for slow (blue) and fast (red) spindles. One asterisk indicates p<0.05, three p<0.001, Wilcoxon signed rank test, n = 58.

We next investigated the spindle-related modulation of CA1 units to the hippocampal ripples. 36% and 33% of the ripples co-occurred with slow and fast spindles respectively (ripples occurred between onset and offset of the spindles). For both slow and fast spindles, the duration of spindle-coupled ripples was larger than the isolated rippled and the duration of slow spindle-coupled ripples was also larger than the duration of fast-spindle coupled ripples (Figure 3A, *x*^2^ = 193.41, d.f. = 2, p = 1.47 × 10^−42^, Kruskall-Wallis test with post hoc Dunn’s multiple correction, p = 3.41 × 10^−5^ for the comparison between slow spindle-coupled ripples and isolated rippled, p = 1.32 × 10^−5^ for the comparison between fast spindle-coupled ripples and isolated rippled, p = 4.11 × 10^−4^ for the comparison between slow spindle-coupled ripples and fast spindle-coupled ripples, mean ± SD, slow spindle-coupled ripples: duration = 0.05 ± 0.02 s, n = 6765, fast spindle-coupled ripples: duration = 0.04 ± 0.01 s, n = 6201, isolated rippled: duration = 0.03 ± 0.02 s, n = 1202). However the amplitude of the isolated ripples was larger than the slow/fast spindle-coupled ripples and the amplitude was larger for fast spindle-coupled ripples than the slow spindle-coupled ripples (Figure 3A, *x*^2^ = 556.9, d.f. = 2, p = 2.23 × 10^−50^, Kruskall-Wallis test with post hoc Dunn’s multiple correction, p = 4.48 × 10^−5^ for the comparison between slow spindle-coupled ripples and isolated rippled; p = 2.21 × 10^−5^ for the comparison between fast spindle-coupled ripples and isolated rippled; p = 3.63 × 10^−4^ for the comparison between slow spindle-coupled ripples and fast spindle-coupled ripples, mean ± SD, slow spindle-coupled ripples: amplitude = 4.92 ± 2.31 SD, n = 6765, fast spindle-coupled ripples: amplitude = 5.49 ± 2.56 SD, n = 6201, isolated rippled: amplitude = 6.32 ± 2.93 SD, n = 1202). Figure 3B displays the distribution of the spindle phases at the time of the ripples (ripple peak) for the ripples co-occurring with spindles obtained by using Hilbert transform. The phases were non-uniformly distributed for both slow and fast spindles so that the majority of the ripples occurred around spindle peak (slow spindle: p < 10^−100^, z = 350.99, Rayleigh test, mean ± SD, phase = − 26.87° ± 36.73°, fast spindle: p < 10^−100^, z = 342.25, Rayleigh test, phase = − 34.64° ± 43.23°, n = 58) corresponding to the active phase of spindles with dominant AD unit activity (Figure 1F). We next compared the modulation of the CA1 units with the ripples for isolated, slow spindle-coupled and fast spindle-coupled ripples (Figure 3C-3E). The modulation index was larger for ripples occurring within the thalamic spindles than the isolated ripples and it was larger for ripples coupled to slow than fast spindles (Figure 3F, *x*^2^ = 116.00, d.f. = 2, p = 6.47 × 10^−26^, Friedman test with post hoc Wilcoxon signed rank test, p = 2.62 × 10^−14^ for the comparison between slow spindle-coupled ripples and isolated rippled, p = 1.14 × 10^−14^ for the comparison between fast spindle-coupled ripples and isolated rippled, p = 2.51 × 10^−13^ for the comparison between slow spindle-coupled ripples and fast spindle-coupled ripples, mean ± SD, slow spindle-coupled ripples: MI = 8.70 ± 0.09 z-score, fast spindle-coupled ripples: MI = 7.90 ± 0.09 z-score, isolated ripples: MI = 5.90 ± 0.08 z-score, n = 58).

**Figure 3.**
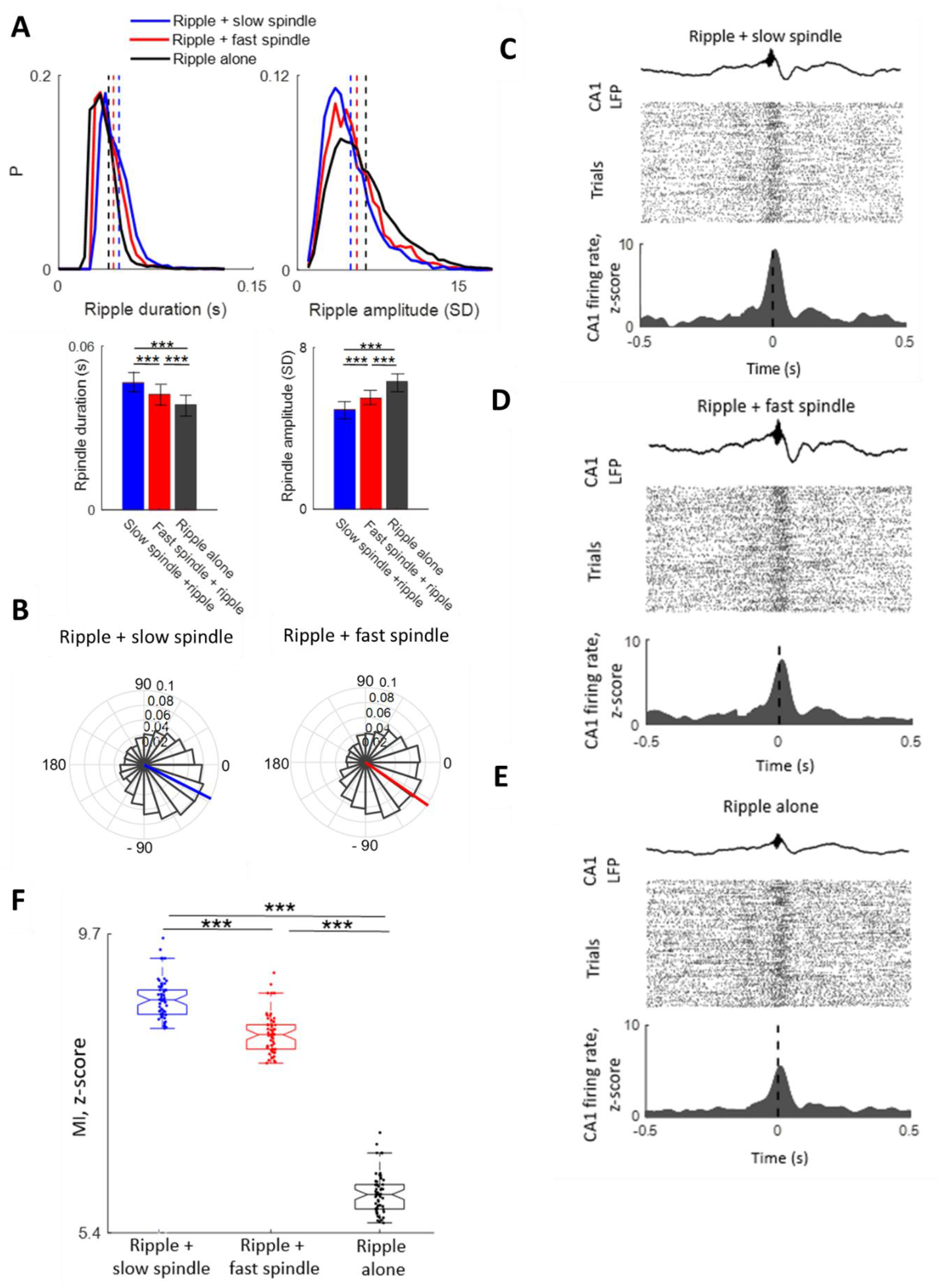
The modulation of CA1 units to the ripples was spindle dependent. (A) Top, histogram of the duration (Left) and amplitude (Right) of ripples co-occurring with slow spindles (blue, ripples occurred between the onset and offset of the spindle), fast spindles (red) and isolated ripples (black). The vertical lines show the mean of the distributions. Bottom, comparison of the duration (Left) and amplitude (Right) of ripples co-occurring with slow spindle (blue), ripples co-occurring with fast spindle (red) and isolated ripples (black, *x*^2^ = 193.41, d.f. = 2, p = 1.47 × 10^−42^, Kruskall-Wallis test with post hoc Dunn’s multiple correction for duration and *x*^2^ = 556.9, d.f. = 2, p =2.23 × 10^−50^, Kruskall-Wallis test with post hoc Dunn’s multiple correction for amplitude, n = 6765, 6201 and 1202 for ripples co-occurring with slow spindle, ripples co-occurring with fast spindle and isolated ripples respectively). (B) Circular histogram of slow (Left) and fast (Right) spindle phases at the time of the ripples (ripple peak) for the ripples co-occurring with spindles obtained by Hilbert transform. (C) Top, representative average raw CA1 LFP traces aligned to peaks of the ripples co-occurring with thalamic slow spindles. Spike rasters (Middle) and peri-ripple-peak histograms (Bottom) of CA1 spikes from one unit (z-score normalized). (D) and (E) The same as (C) for ripples co-occurring with thalamic fast spindles and isolated ripples respectively. (F) Comparison between the modulation index of CA1 units for ripples co-occurring with slow spindles (blue), fast spindles (red) and isolated ripples (*x*^2^ = 116.00, d.f. = 2, p = 6.47 × 10^−26^, Friedman test with post hoc Wilcoxon signed rank test, n = 58). Three asterisk indicates p<0.001.

In sum, we found that the modulation of CA1 units with spindles and the spindle-associated modulation of CA1 units with ripples were stronger for slow than fast spindles. These results suggest that spindles with lower frequencies provide a wider depolarizing window for the temporal coordination of the hippocampal and thalamocortical networks and enhancement of the multi-regional interactions.

### Stronger modulation of hippocampal ripples to longer thalamic spindles

We next explored the temporal association between hippocampal ripples and thalamic spindles. First the event correlation histograms of ripple events time-locked to the spindle onset were computed which revealed increase of ripple occurrence after the spindle onset with a peak of around 200 ms for both slow and fast spindles (Figure4-Figure supplement 1). To further explore the coupling between spindles and ripples, we next quantified the modulation of hippocampal ripple amplitude with the phase of slow/fast spindles detected from the thalamic LFP using phase amplitude coupling (PAC) analyses (Figure 4A-4C, see Materials and methods for details). The ripples were mostly coupled to the phases around the peak of thalamic spindles with the SI angles dominantly after the spindle peak for the slow spindles and before the spindle peak for the fast spindles (slow spindle: p = 1.81 × 10^−3^, z = 1805.4, Rayleigh test, mean ± SD, SI angle = 29.72° ± 60.17°, n = 190006, fast spindle: p = 1.61 × 10^−3^, z = 1098.3, Rayleigh test, SI angle = − 35.67° ± 64.23°, n = 180175). In addition, the ripples were more strongly coupled to the slow than fast spindles (Figure 4D, p = 2.13 × 10^−5^, Mann-Whitney U-test, mean ± SD, slow spindle: SI strength = 0.69 ± 0.23, fast spindle: SI strength = 0.54 ± 0.22).

**Figure 4.**
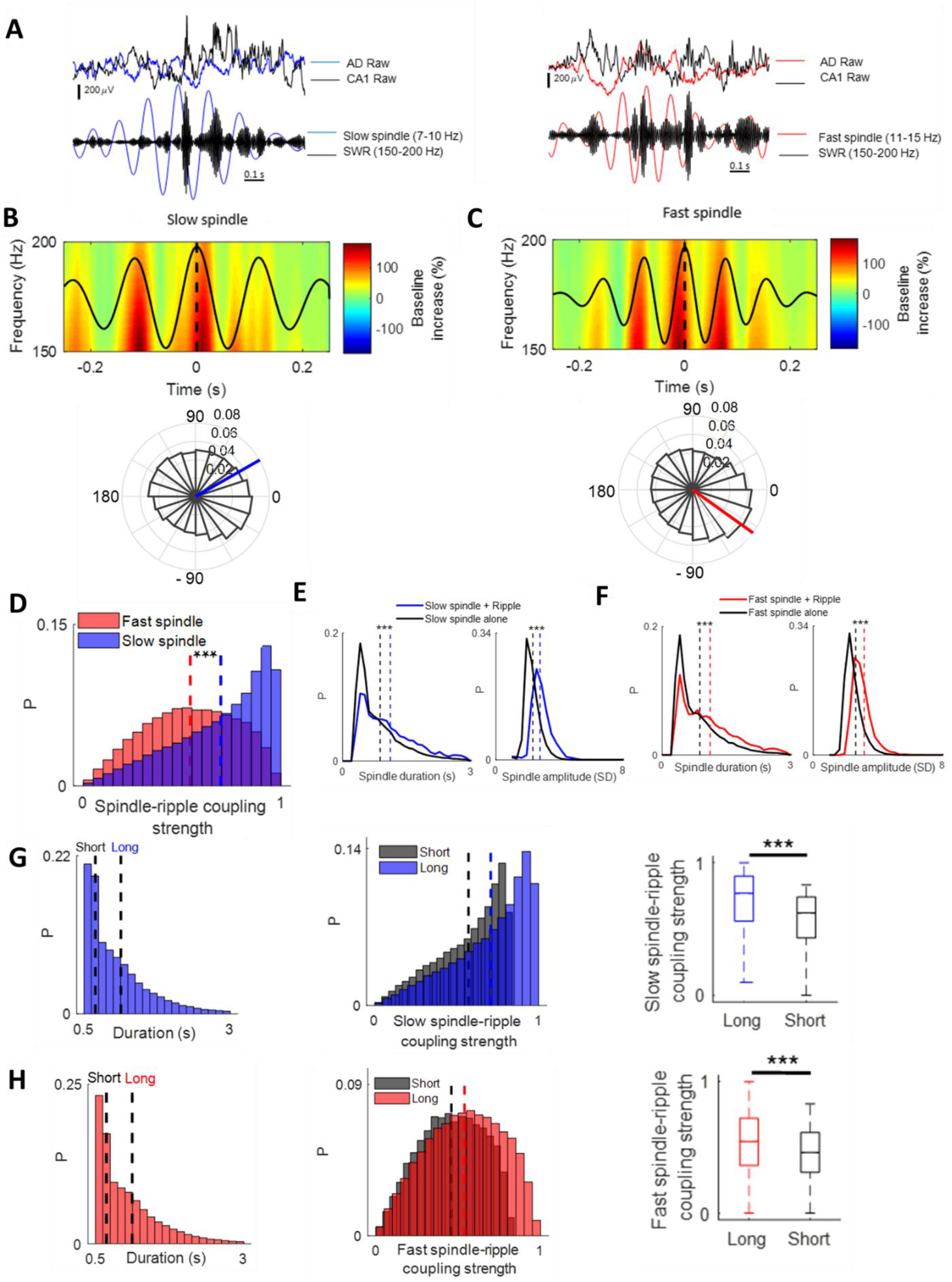
Temporal coordination of slow and fast spindles with ripples. (A) Raw (Top) and filtered (Bottom) traces of AD and CA1 (black) LFPs during one slow (Left, blue) and fast (Right, red) spindle co-occurring with ripple events. The AD and CA1 LFPs were filtered in the spindle (slow spindle: 7-10 Hz, fast spindle: 11-15 Hz) and ripple (150-200 Hz) frequency ranges respectively. (B) Top, average of slow spindle peak–locked TFR of the CA1 LFP (percentage change from pre-event baseline, n= 190006). Black curves show grand average filtered thalamic LFP in the slow spindle frequency range (7-10 Hz) aligned to the spindle peak (time 0). Bottom, circular histogram of SI angles of the ripples relative to the slow spindles. SI angles were non-uniformly distributed (p = 1.81 × 10^−3^, z = 1805.46, Rayleigh test). (C) The same as (B) for fast spindles (11-15 Hz, p = 1.61 × 10^−3^, z = 1098.3, Rayleigh test). (D) Histogram of the spindle– ripple coupling strength for slow (blue) and fast (red) spindles. (E) Histogram of the duration (Left) and amplitude (Right) of slow spindles co-occurring with ripples (blue) and isolated spindles (black). (F) The same as (E) for fast spindles. (G) Histogram of slow spindle duration with the cut-offs (vertical lines) for short (mean ± SD: duration = 0.52 ± 0.06 s, n= 63335) and long slow spindle events (mean ± SD: duration =1.81 ± 0.53 s) defined as first and third tertiles respectively (Left). The distribution of the slow spindle-ripple coupling strength for short (black) and long (blue) slow spindles (Middle). The vertical lines show the average of the distributions. Comparison of the slow spindle-ripple coupling strength for short (black) and long (blue) slow spindles (Right). (H) The same as (G) for fast spindle (mean ± SD: duration = 0.51 ± 0.06 s and 1.78 ± 0.53 s for short and long fast spindles respectively, n= 60058). Three asterisk indicates p<0.001, Mann-Whitney U-test. **Figure supplement 1.** The increase of ripple occurrence after the spindle onset.

We next compared the duration and amplitude of spindles co-occurring with ripples (spindles with at least one ripple between their onset and offset) with isolated spindles. 28.49% and 27.29% of the slow and fast spindles were coupled to the ripples. The duration and amplitude of both slow and fast spindles co-occurring with ripples were larger than the isolated spindles (Figure 4E-4F, slow spindle: p < 10^−100^ for both duration and amplitude, Mann-Whitney U-test, mean ± SD, duration = 1.30 ± 0.62 s and amplitude = 2.77 ± 0.98 SD, n = 54151 for slow spindles coupled with ripples, duration = 1.02 ± 0.52 s and amplitude = 2.21 ± 0.7 SD, n = 135855 for isolated slow spindles, fast spindle: p < 10^−100^ for both duration and amplitude, Mann-Whitney U-test, duration = 1.31 ± 0.63 s and amplitude = 3.14 ± 0.82 SD, n = 49187 for fast spindles coupled with ripples, duration = 1.01 ± 0.51 s and amplitude = 2.62 ± 0.7 SD, n = 130988 for isolated fast spindles).

We further explored whether the spindle-ripple coupling depended on the duration of the spindles. To assess this possibility, we partitioned the distribution of the spindle durations into tertiles and compared the strength of the coupling of ripples to long (third tertile) versus short (first tertile) spindles. For both slow and fast spindles, the spindle-ripple coupling strength was larger for long versus short spindles (Figure 4G-4H, slow spindle: p = 3.02× 10^−4^, Mann-Whitney U-test, mean ± SD, SI strength = 0.70 ± 0.22 for long and 0.57 ± 0.19 for short slow spindles, fast spindle: p = 1.09× 10^−3^, Mann-Whitney U-test, SI strength = 0.53 ± 0.22 for long and 0.45 ± 0.19 for short fast spindles). These results indicate the stronger coupling of ripples to slow than fast spindles and the enhancement of inter-regional communications during spindles with longer duration.

### Spindle occurring closer to the SO trough were more strongly coupled to the ripples

We next investigated the temporal interaction of SOs and ripples/spindles. 53.79% of the ripples co-occurred with SOs (SO peak occurred within ± 0.5 s interval around ripple peak). The duration of the ripples co-occurring with SOs was larger compared to the isolated ripples while the amplitude of these ripples was lower than the ripples occurring alone (Figure5-Figure supplement 1A, p = 1.35×10^−75^ and p = 6.46×10^−10^ for duration and amplitude respectively, Mann-Whitney U-test, mean ± SD, duration = 0.04 ± 0.009 s and amplitude = 6.11 ± 4.20 SD, n = 10110 for ripples coupled with SOs, duration = 0.03 ± 0.008 s and amplitude = 6.36 ± 3.10 SD, n = 8683 for isolated ripples). In addition, the phases of SO at the time of ripple events (ripple peak) were highly non-uniformly distributed (p < 10^−100^, z =674.04, Rayleigh test, n = 18793) with a mean phase of 152.4 ° ± 80.43° corresponding to more dominant ripple activity before SO trough (Figure5-Figure supplement 1B).

79.83% and 79.78 % of the slow and fast spindles co-occurred with SOs respectively (SO peak occurred within ± 1.5 s interval around spindle maximum peak). The duration and amplitude of the slow/fast spindles co-occurring with SOs were larger compared to the isolated spindles (Figure5-Figure supplement 2, slow spindle: p = 5.10×10^−82^ and p = 1.53×10^−72^ for duration and amplitude respectively, Mann-Whitney U-test, mean ± SD, duration = 1.05 ± 0.62 s and amplitude = 2.89 ± 0.85 SD, n = 151700 for slow spindles coupled to SOs, duration = 0.88 ± 0.52 s and amplitude = 2.51 ± 0.73 SD, n = 38306 for isolated slow spindles, fast spindle: p = 9.92×10^−58^ and p = 5.15×10^−96^ for duration and amplitude respectively, Mann-Whitney U-test, duration = 1.05 ± 0.62 s and amplitude = 3.14 ± 0.88 SD, n = 143761 for fast spindles coupled to SOs, duration = 0.87 ± 0.51 s and amplitude = 2.72 ± 0.74 SD, n = 36414 for isolated fast spindles). Figure 5A displays the circular histogram of the SO phases (using Hilbert transform) at the time of the slow and fast spindle events (slow spindle: p = 5.10×10^−52^, z = 2447.9, Rayleigh test, mean ± SD, phase = 140.06 ° ± 78.80°, n = 190006, fast spindle: p = 4.71×10^−42^, z = 2512.4, Rayleigh test, phase = 121.69 ° ± 80.81°, n = 180175). We hypothesized that the SO phase at the time of the spindle occurrence predicts the strength of the coupling of the ripples to the spindle (Figure 5B). Hence we next computed the circular-linear correlation between the phase of SO at the time of the spindle and the spindle-ripple coupling strength (obtained from PAC analyses) separately for slow and fast spindles. Interestingly, the locking phase of spindles to SOs was highly predictive of the spindle-ripple coupling strength especially for slow spindles so that spindles that occurred closer to SO trough corresponding to depolarized phase of the SO were more strongly coupled to the ripples (Figure 5C-5D, slow spindle: p = 3.70×10^−14^, r = 0.79, circular-liner correlation, n = 190006, fast spindle: p = 2.79×10^−6^, r = 0.51, circular-liner correlation, n = 180175). These results further suggest the multi-regional communications at the time of the spindles and are consistent with a previous study showing the increase of spindle-ripple coupling as a result of spindle-like stimulation in-phase with slow oscillation up-states (Latchoumane et al., 2017).

**Figure 5.**
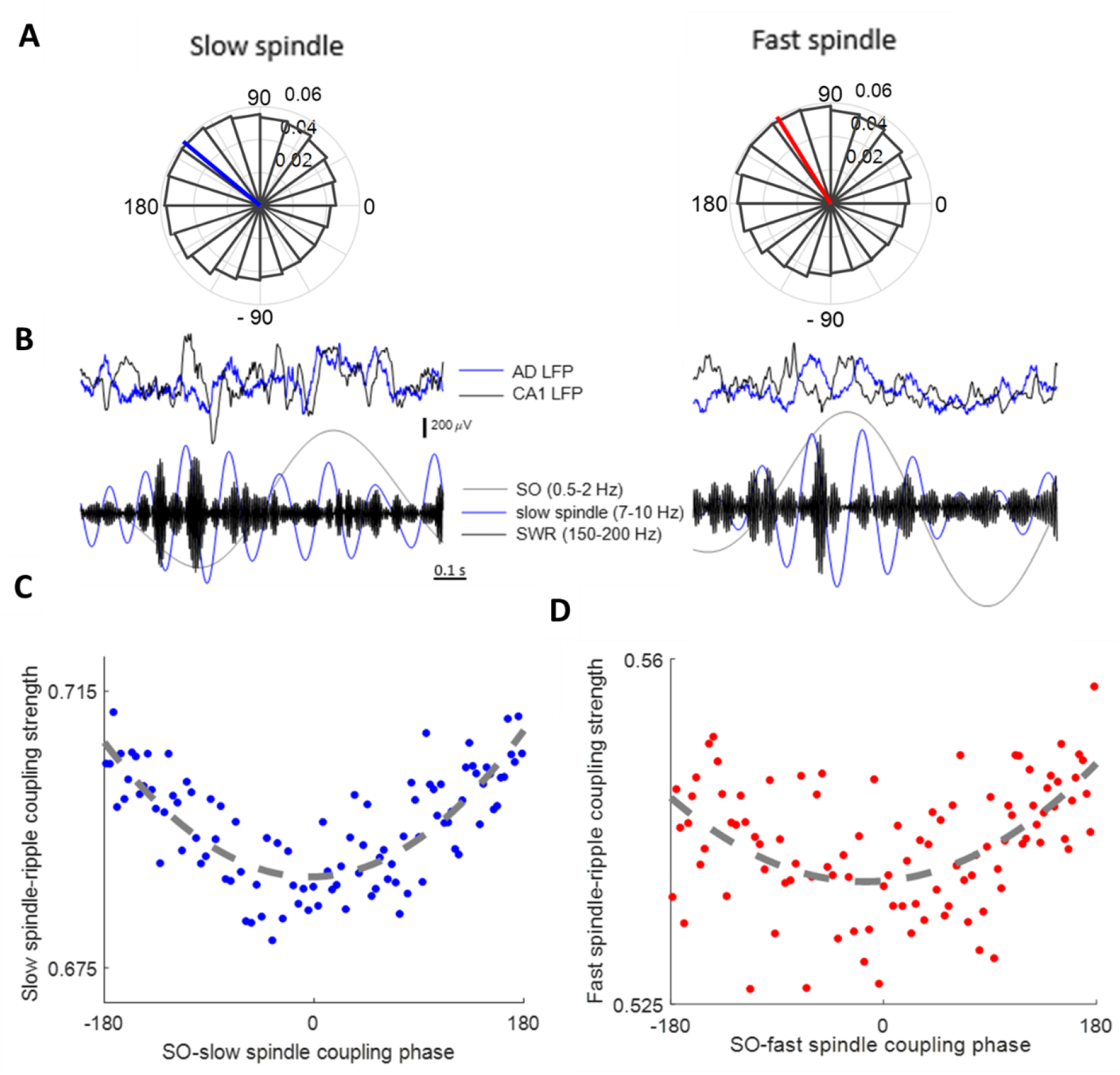
SO-spindle coupling phase predicts spindle-ripple coupling strength. (A) The circular histogram of the SO phases at the time of slow (Left) and fast (Right) spindles (spindle peak). (B) Raw (Top) and filtered (Bottom) traces of AD (blue) and CA1 (black) LFPs during two slow spindles occurring around trough (Left) and peak (Right) of the SO showing stronger spindle-ripple coupling for the spindle occurring around SO trough. The AD and CA1 LFPs were filtered in the SO/spindle (SO: 0.5-2 Hz, slow spindle: 7-10 Hz) and ripple (150-200 Hz) frequency ranges respectively. (C) The slow spindle-ripple coupling strength versus the SO phase at the time of the slow spindle (100 bins). The dotted gray line shows the quadratic fit to the data indicating of large circular-linear correlation. (D) The same as (C) for fast spindles. **Figure supplement 1.** Temporal associations between SOs and ripples. **Figure supplement 2**. Differential properties of spindles co-occurring with SOs compared to the isolated spindles.

### Stronger coupling of CA1 units to thalamic spindles after exploration

We next examined whether the temporal association of CA1 units/ripple and thalamic spindles was task-relevant. To this end, we computed the spindle/ripple duration, spindle-associated modulation of CA1 units and spindle-ripple coupling during sleep periods that preceded and followed spatial exploration. The duration of both slow and fast spindles as well as the ripples were larger for sleep after than before the exploration (Figure 6A, slow spindle: p = 3.08×10^−5^, Mann-Whitney U-test, mean ± SD, duration = 1.02 ± 0.51 s, n = 62991 for after exploration and 0.85 ± 0.42 s, n = 52968 for before exploration, fast spindle: p = 2.04×10^−4^, Mann-Whitney U-test, duration = 1.01 ± 0.5 s, n = 56518 for after exploration and 0.84 ± 0.42 s, n = 47514 for before exploration, ripple: p = 1.95×10^−4^, Mann-Whitney U-test, duration = 0.04 ± 0.03 s, n = 7588 for after exploration and 0.03 ± 0.02 s, n = 7485 for before exploration). During the sleep periods following the exploration, CA1 units were more strongly modulated by both slow and fast spindles compared to the sleep before exploration (Figure 6B-6D, *x*^2^ = 174.00, d.f. = 1, p = 7.24 × 10^−16^, Friedman test with post hoc Wilcoxon signed rank test, p = 2.23 ×10^−14^ for comparison between the modulation index of CA1 units with slow spindle before and after exploration, p = 3.64 ×10^−13^ for comparison between the modulation index of CA1 units with fast spindle before and after exploration, p = 1.51 ×10^−14^ for comparison between the modulation index of CA1 units with slow and fast spindle after exploration, p = 1.11 ×10^−14^ for comparison between the modulation index of CA1 units with slow and fast spindle before exploration, mean ± SD, slow spindle: MI = 4.71 ± 0.52 z-score for after and 3.11 ± 0.51 z-score for before exploration, fast spindle: MI = 4.01 ± 0.51 z-score for after and 2.51 ± 0.52 z-score for before exploration, n = 58). We next compared the phase-locking of CA1 unit spiking to slow/fast spindles before and after exploration (Figure 6E). While the spikes of CA1 units mostly occurred before the slow spindle peak during pre-sleep periods, they occurred at more positive phases of slow spindles with a mean phase slightly after the slow spindle peak (slow spindle: p = 1.62 × 10^−4^, circular-mtest, mean ± SD, phase = 17.71 ° ± 92.81° for after and −11.71 ° ± 95.80° for before exploration, fast spindle: p = 0.1, circular-mtest, phase = −23.53 ° ± 82.51° for after and −24.69 ° ± 86.8° for before exploration, n = 58).

**Figure 6.**
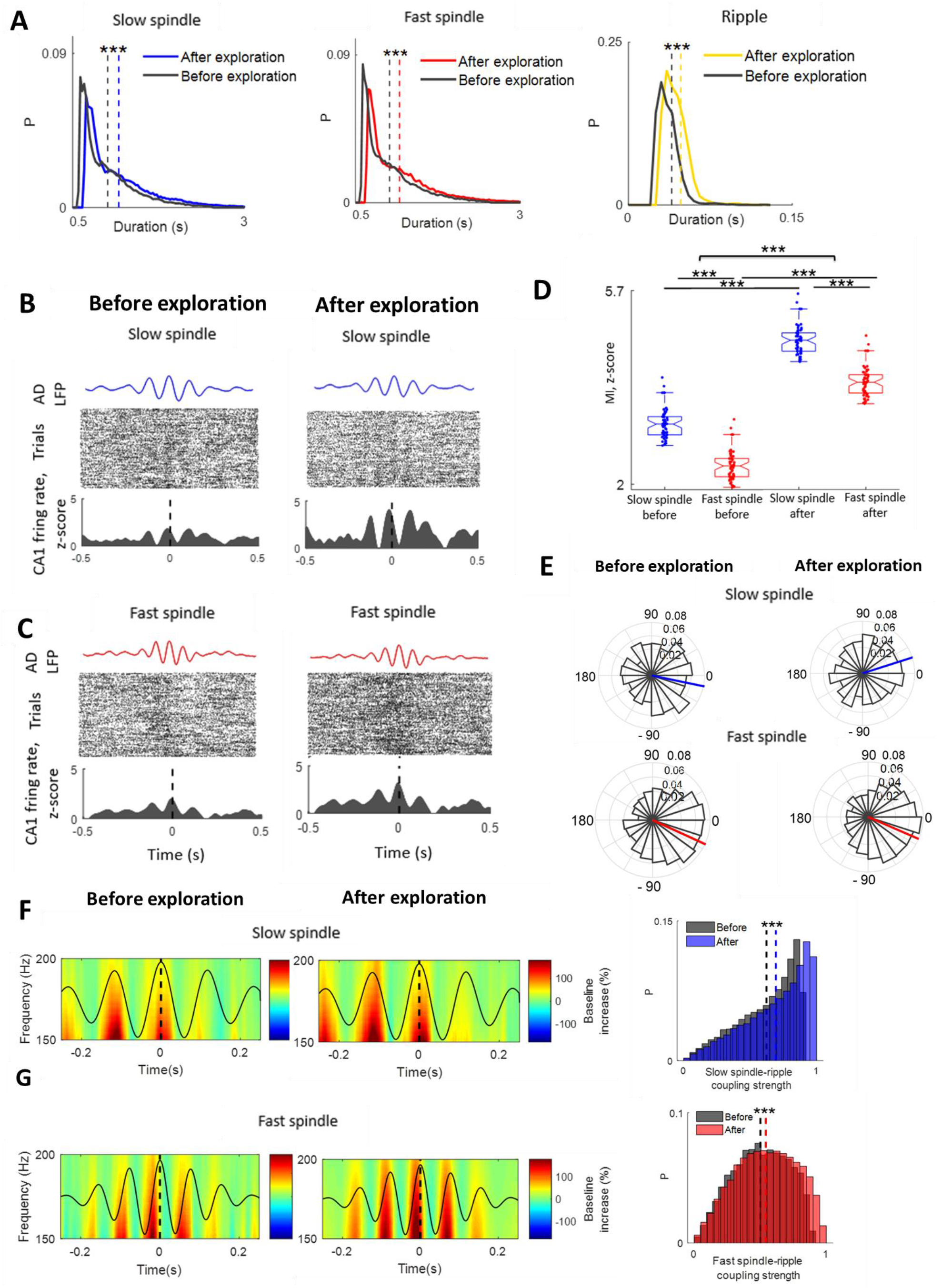
Modulation of CA1 units and ripples to thalamic spindles enhance after exploration. (A) Histogram of the duration of slow spindles (Left), fast spindles (Middle) and ripples (Right) for before (black) and after (blue, red, yellow for slow spindles, fast spindles and ripples respectively) exploration. (B) Top, representative average AD LFP traces filtered in the slow spindle range (7-10 Hz) aligned to the slow spindle peaks. Spike rasters (Middle) and peri-slow spindle-peak histograms (Bottom) of CA1 spikes from one unit (z-score normalized) before (Left) and after (Right) exploration. (C) The same as (B) for fast spindles (11-15 Hz). (D) Comparison of the modulation index of CA1 units with slow (blue) and fast (red) spindles before and after exploration (*x*^2^ = 174.00, d.f. = 1, p = 7.24 × 10^−16^, Friedman test with post hoc Wilcoxon signed rank test, n = 58). (E) Circular histograms of slow (Top) and fast (Bottom) spindle phases at the time of CA1 unit spikes before (Left) and after (Right) exploration. (F) Average of slow spindle peak–locked TFR of the CA1 LFP (percentage change from pre-event baseline) before (Left, n =52968) and after (Middle, n= 62991) exploration. Black curve shows grand average filtered thalamic LFP in the slow spindle frequency range (7-10 Hz) aligned to the spindle peak (time 0). Histograms of the slow spindle-ripple coupling strength before (black) and after (blue) exploration (Right). (G) The same as (F) for fast spindles (11-15 Hz). Three asterisk indicates p<0.001.

In addition, PAC analyses revealed that the spindle-ripple coupling strength was larger during sleep after than before exploration for both slow and fast spindles (Figure 6F-6G, slow spindle: p = 2.08×10^−8^, Mann-Whitney U-test, mean ± SD, SI strength = 0.69 ± 0.23, n= 62991 for after exploration and 0.62 ± 0.21, n = 52968 for before exploration, fast spindle: p = 1.39×10^−9^, Mann-Whitney U-test, SI strength = 0.54 ± 0.22, n = 56518 for after exploration and 0.50 ± 0.2, n = 47514 for before exploration). These results reveal task-specific interactions between the hippocampal and thalamic activity during spindles/ripples suggesting functional role of thalamic spindles on cross regional information transfer.

### Hippocampal-thalamocortical model for multi-regional interactions during spindles

We next developed a simplified hippocampal-thalamocortical neural mass model to investigate the mechanisms underlying the hippocampal-thalamocortical temporal interactions during thalamic spindles. The model consists of thalamocortical and CA1-CA3 neural networks with long-range bidirectional hippocampal-thalamocortical and cortico-thalamic projections. As we previously described (Ghorbani et al., 2012; Hashemi et al., 2019; Azimi et al., 2021), in this model the up and down states with high frequency activity during the up states are generated in recurrently connected cortical networks due to dendritic spike frequency adaptation. Similar to previous studies (Cona et al., 2014; Destexhe et al., 1996; Destexhe and Sejnowski, 2003; Hashemi et al., 2019) spindles are generated in a bursting recurrent thalamic network which in our model simply consists of one identical group of inhibitory neurons representing the thalamic reticular neurons and one identical group of thalamocortical neurons. The ripples are produced due to interactions between CA1 excitatory and inhibitory neurons. While the model is deterministic (with no external or intrinsic noise), due to nonlinear interaction of multiple thalamocortical and hippocampal oscillators, it can generate spindles and ripples with variable frequency, duration and amplitude by chaotic dynamics (Figure 7A, Ghorbani et al., 2012; Hashemi et al., 2019).

**Figure 7.**
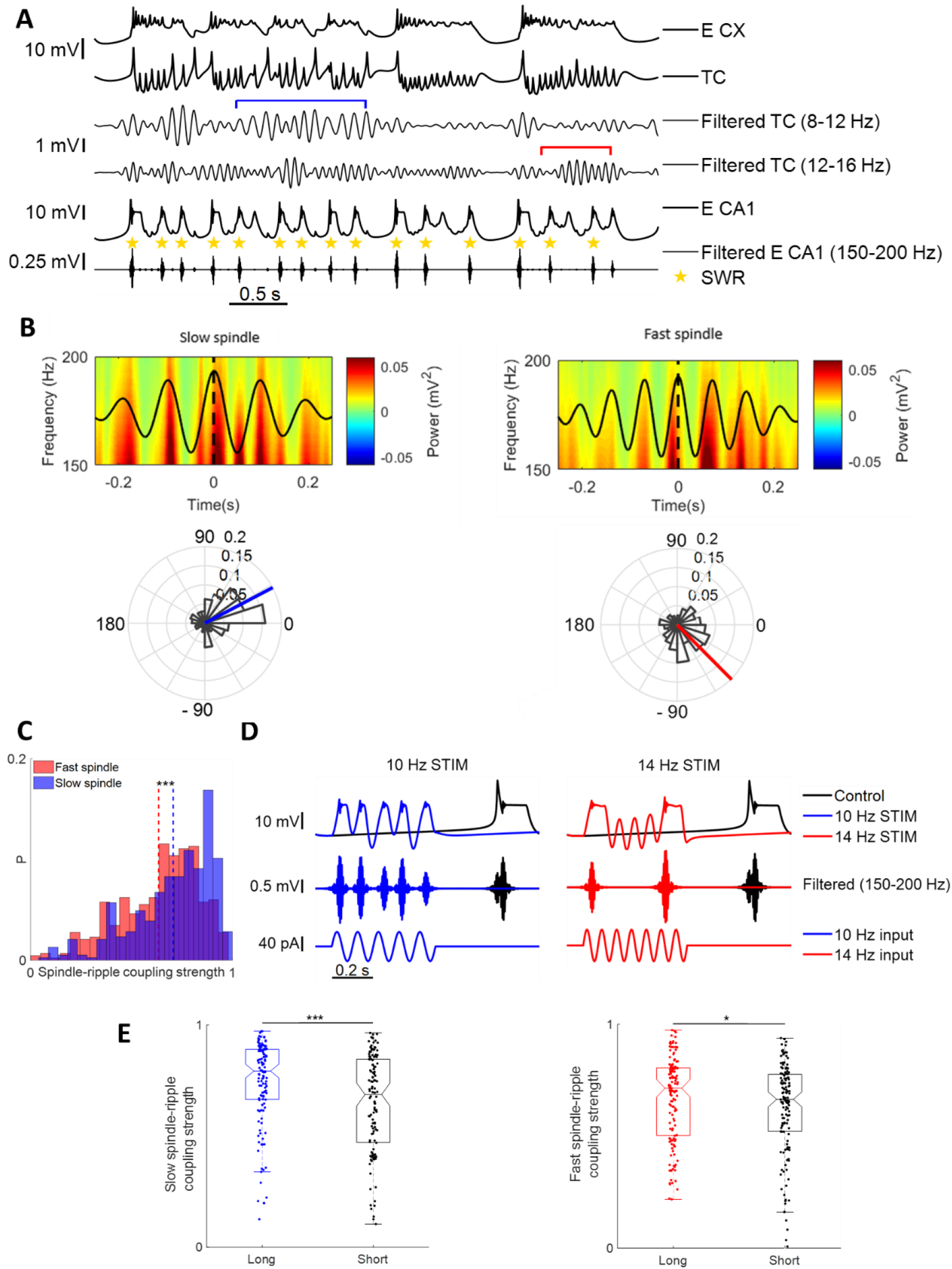
Hippocampal-thalamocortical temporal interactions during spindles in the model. (A) From top to bottom respectively: Broadband trace of the membrane potential of excitatory neurons of the cortical network, broadband trace of the membrane potential of thalamocortical neurons, filtered (8-12 Hz) trace of the membrane potential of thalamocortical neurons with detected slow spindles (blue bracket), filtered (12-16 Hz) trace of the membrane potential of thalamocortical neurons with detected fast spindles (red bracket), broadband trace of the membrane potential of excitatory neurons of CA1 network and filtered (150-200 Hz) traces of the membrane potential of CA1 excitatory neurons with detected ripples (yellow stars). (B) Left: Top, average of slow spindle peak–locked TFR of the CA1 excitatory neuron membrane potential (n= 385). Black curve shows grand average filtered thalamocortical neuron membrane potential in the slow spindle frequency range (8-12 Hz) aligned to the spindle peak (time 0). Bottom, circular histogram of SI angles of the ripples relative to the slow spindles. SI angles were non-uniformly distributed (p = 2.50 × 10^−30^, z = 65.23, Rayleigh test). Right: The same as Left for fast spindles (12-16 Hz, p = 2.75 × 10^−16^, z = 35.13, Rayleigh test). (C) Histogram of the spindle–ripple coupling strength for slow (blue) and fast (red) spindles. (D) The effect of applying input currents (Bottom) oscillating in the slow spindle (Left, 10 Hz) and fast spindle (Right, 14 Hz) frequency ranges on the trace of CA1 excitatory neurons when the thalamocortical network was removed. The stimulation was applied during ripple refractory time. The control and stimulated raw (Top) and filtered (150-200 Hz) traces (Middle) are shown in black, blue (slow spindle)/red (fast spindle) respectively. (E) Comparison of the spindle-ripple coupling strength between short (black) and long (blue) spindles for slow (Left) and fast (Right) spindles. One asterisk indicates p<0.05, three p<0.001, Wilcoxon signed rank test. **Figure supplement 1.** Hippocampal-thalamocortical temporal interactions during spindles in the model data.

To investigate the modulation of the thalamocortical and CA1 neurons with thalamic spindles in the model, we first computed the average firing rate of thalamocortical and CA1 neurons around the thalamic spindle peak separately for slow (8-12 Hz) and fast (12-16 Hz) spindles (Figure7-Figure supplement 1A-B). Similar to the experimental findings, the firing rate of both thalamocortical and CA1 neurons were more strongly modulated by slow than fast spindles. We next quantified the spindle-ripple coupling strength and phase using PAC analyses in the model data (Figure 7B). Consistent with the experimental results, the hippocampal ripples were mostly coupled to the phases around the peak of thalamic spindles with the SI angles dominantly after the spindle peak for the slow spindles and before the spindle peak for fast spindles (slow spindle: p = 2.50 × 10^−30^, z = 65.23, Rayleigh test, mean ± SD, SI angle = 27.78 ± 3.89 °, n = 385, fast spindle: p = 2.75 × 10^−16^, z = 35.13, Rayleigh test, SI angle = − 43.90° ± 4.36 °, n = 435). Spindle-ripple coupling strength was larger for slow than fast spindles indicating stronger coupling of ripple amplitude to the phase of slow spindles (Figure 7C, p = 6.19 × 10^−9^, Mann-Whitney U-test, slow spindle: mean ± SD, SI strength = 0.71 ± 0.20, fast spindle: SI strength = 0.63 ± 0.20).

In order to explore the mechanism underlying the stronger coupling of ripples to slow versus fast spindles, we removed the thalamocortical projections to the hippocampal network and compared the effect of applying transient input currents oscillating at the slow versus fast spindle frequency ranges (0.5 s, 85 pA peak to trough). As is shown in figure 7D, while 10 Hz stimulation induced ripple activity at each of the input current cycles at around the peak of oscillations, 14 Hz stimulation could not provide the large enough temporal window at each cycle required for ripple generation in the CA1 network because of lower duration of the active and quiescence phases of the input current. These results explain the reason behind stronger coupling of slow versus fast spindles to the hippocampal ripples indicating the role of spindles with slower frequencies on organizing the ripple dynamics and facilitating multi-regional temporal coordination required for memory consolidation.

In agreement with the experimental results (Figure 4E-4F), in the model data, the duration and amplitude of both slow and fast spindles co-occurring with ripples were larger than the isolated spindles (Figure7-Figure supplement 1C-D, slow spindle: p = 6.70 × 10^−3^ and p = 4.83× 10^−6^ for duration and amplitude respectively, Mann-Whitney U-test, mean ± SD, duration = 0.76 ± 0.24 s and amplitude = 2.85 ± 0.60 SD, n = 367 for slow spindles coupled with ripples, duration = 0.63 ± 0.13 s and amplitude = 2.12 ± 0.50 SD, n = 18 for isolated slow spindles, fast spindle: p = 3.80 × 10^−2^ and p = 2.80 × 10^−2^ for duration and amplitude respectively, Mann-Whitney U-test, mean ± SD, duration = 0.82 ± 0.23 s and amplitude = 3.25 ± 0.45 SD, n = 411 for fast spindles coupled with ripples, duration = 0.72 ± 0.15 s and amplitude = 3.05 ± 0.42 SD, n = 24 for isolated fast spindles).

To assess whether the spindle-ripple coupling depended on the duration of the spindles in the model data, we next partitioned the distribution of the spindle durations into tertiles and compared the coupling strength of ripples to long (third tertile) versus short (first tertile) spindles. Consistent with our experimental findings, the ripples were more strongly coupled to long versus short spindles for both slow and fast spindles (Figure 7E, slow spindle: p = 2.56 × 10^−4^, Mann-Whitney U-test, mean ± SD, SI strength = 0.74 ± 0.19 for long and 0.65 ± 0.22 for short slow spindles, fast spindle: p = 4.03 × 10^−2^, Mann-Whitney U-test, SI strength = 0.67 ± 0.19 for long and 0.62 ± 0.20 for short fast spindles). These results further support our hypothesis that the spindles with slower oscillating frequency and longer duration enhance the hippocampal-thalamocortical interaction by providing a longer window of opportunity for synchronization.

### Figure 7—source code 1

Matlab code for generating the model data

## Discussion

Together, our results provide the first demonstration that spindle frequency and duration can provide valuable information about the underlying multi-regional interactions essential for memory consolidation computations. Our findings indicate enhanced thalamic spindle duration and spindle-associated activity of CA1 units during post-learning sleep. The other key finding revealed by the current study is that the modulation of the CA1 units to the hippocampal ripples is thalamic activity dependent so that the depolarizing window during the thalamic spindle peaks provide the fine-tuned window for CA1 activity coordination. We further described how the low frequency long-duration spindles can increase the temporal multi-regional coordination by developing a simplified hippocampal- thalamocortical model.

Several studies have inferred the phase locking of hippocampal ripples to neocortical (Mölle et al., 2006; Ngo et al., 2020; Siapas and Wilson, 1998; Sirota et al., 2003; Varela and Wilson, 2020; Peyrache et al., 2011) or hippocampal/parahippocampal (Clemens et al., 2011; Helfrich et al., 2019; Jiang et al., 2019; Staresina et al., 2015; Ngo et al., 2020) spindles. Manipulating studies indicated the role of spindle-ripple coupling in successful memory consolidation (Binder et al., 2019; Latchoumane et al., 2017; Maingret et al., 2016; Xia et al., 2017). The modulation of the CA1 unit activity with cortical spindles has been shown previously (Sirota et al., 2003; Varela and Wilson, 2020; Sullivan et al., 2014; Wierzynski et al., 2009). A recent study explored the effect of before/after exploration on the modulation of CA1 units with cortical spindles. Although the modulation of most of CA1 units with spindles increased after exploration the difference was not found to be significant. However none of these previous studies used thalamic spindles to explore the precise temporal coupling of hippocampal activity to spindles which are originated in the thalamus and can reach the hippocampus by direct projections from the thalamus to CA1 (Vertes, 2015). In addition how the hippocampal-thalamocortical interactions during spindles depend on spindle properties was not investigated in the previous studies. We sought to address this gap by focusing on thalamic spindles to investigate the differential contribution of spindles with different duration and oscillating frequency to multi-regional coordination. We particularly showed that the modulation of CA1 unit activity with thalamic spindles as well as spindle-ripple coupling strength enhanced after exploration. The repeated activation of CA1 units during thalamic spindle peaks which provide fine-grained temporal windows of synaptic excitability can facilitate spike-timing-dependent synaptic plasticity. We further explained how spindles with slower frequencies can enhance the hippocampal-thalamocortical temporal interactions by providing temporally large enough successive active and quiescent windows. Consistently reports have shown association between an increase in spindle frequency during NREM and memory impairments in old subjects (Taillard et al., 2019) and negative correlations between spindle frequency and cognitive abilities in children (Chatburn et al., 2013; Geiger et al., 2011). However the significance of spindle frequency in rodents remains to be elucidated. A recent study suggested that memory impairment in a hippocampus-dependent object place recognition task in young sleep deprived mice could be associated with an increase in spindle frequency (Yuan et al., 2021).

The functional relevance of long-duration ripples for successful memory consolidation has been underscored recently (Fernández-Ruiz et al., 2019; Ngo et al., 2020). Consistently we found that the ripple duration increased after exploration. We further found an increase in the spindle duration post-behavior and provided evidence for the importance of long-duration spindles for hippocampal-thalamocortical interactions. Increase in spindle duration in the sleep following learning was reported in humans (Morin et al., 2008; Fogel et al., 2007). Several neuropsychiatric disorders are also associated with reduction of spindle duration (Ferrarelli et al., 2007; Fernandez and Lüthi, 2020) and spindle duration also decreases with age (Nicolas et al., 2001). In accordance with these studies, our findings highlight the significance of spindle duration for successful memory consolidation which needs to be explored more directly in the future manipulating studies. While hippocampal ripples are nested in cortical spindle troughs (Varela and Wilson, 2020; Binder et al., 2019; Oyanedel et al., 2020), we found that the ripples mostly occurred at peaks of thalamic spindles. The reason behind this differential phase-locking of ripples to cortical versus thalamic spindles is that the thalamic units are active at peaks and silent at troughs of the thalamic spindles while cortical units show an inverse spike-field coupling with higher activity at troughs of cortical spindles (Bandarabadi et al., 2020). A recent study reported increase in MD multi-unit activity with spindle-coupled ripples and decrease in MD firing rate during isolated ripples (Yang et al., 2019). The responsiveness of cortical units to hippocampal ripples was shown to be greater during spindles rather than outside spindles (Peyrache et al., 2011). Our findings demonstrated that CA1 neurons were highly modulated with ripples co-occurring with thalamic spindles compared to isolated ripples. Interestingly we also found that CA1 units were locked to more delayed phases of slow spindles in the sleep post-behavior so that they were more active after the spindle peaks. These results provide the first evidence for the functionally relevance of the spindle phase at the time of CA1 spikes. The precise temporal coupling of fast spindles to SOs was suggested to be of functional benefit for the processing of memory (Kim et al., 2019; Niknazar et al., 2015; Helfrich et al., 2018; Ladenbauer et al., 2017; Latchoumane et al., 2017). However the functional relevance of SO-slow spindle coupling is unclear (Muehlroth et al., 2019). The enhancement of spindle-ripple coupling and memory consolidation by thalamic fast spindle induction phase-locked to SOs have been reported (Latchoumane et al., 2017). Our results extend these findings showing a large circular-linear correlation between the coupling phase of both thalamic slow and fast spindles to SOs and the strength of spindle-ripple coupling so that phase-locking of spindles closer to SO troughs representing the up state was associated with stronger spindle-ripple coupling. We found both thalamic slow and fast spindles were coupled to SOs occurring mainly during the down to up state transition. These results are consistent with studies reported the coupling of cortical SOs and fast spindles in humans (Mölle et al., 2011; Clemens et al., 2011; Klinzing et al., 2016; Oyanedel et al., 2020; Andrillon et al., 2011; Mak-McCully et al., 2017) or rodents (Mölle et al., 2006; Peyrache et al., 2011; Niethard et al., 2018) but are not in accordance with EEG studies in humans showing the main occurrence of cortical slow spindles during the cortical up to down state transition (Mölle et al., 2011; Clemens et al., 2011; Klinzing et al., 2016). In addition the differential topographical distribution of slow and fast spindles in humans has not been observed in rodents (Kim et al., 2015). Unlike the human EEG/LFP spectrum in which both slow and fast spindle peaks are distinguishable (Dehnavi et al., 2021; Andrillon et al., 2011), the rodents spectrum shows no distinct peak at the spindle frequency range (7–15 Hz). (Mölle et al., 2009) so that distinguishing fast and slow spindles from the spectrum is not possible in rodents. We recently reported that slow spindles are associated with cortical high frequency oscillations so that they can be generated as a result if nonlinear interaction of thalamic and cortical oscillators (Hashemi et al., 2019). The differential properties of fast and slow spindles in humans versus rodents might be due to differences in high frequency activity spatiotemporal distributions.

Computational models showed that the interaction of thalamocortical and reticular neurons in bursting recurrent thalamic networks can generate fast spindles (Cona et al., 2014; Destexhe et al., 1996; Destexhe and Sejnowski, 2003; Hashemi et al., 2019). In our model consistent with previous reports (Bojean et al., 2011), the beginning and termination of spindles are controlled by the long-range cortico-thalamic input (Hashemi et al., 2019). Progressive changes in the activity of thalamocortical (Bal and McCormick, 1996; Lüthi and McCormick, 1998; Lüthi et al., 1998) or reticular neurons (Bal et al., 1995; Kim and McCormick, 1998; Bartho et al., 2014) during the spindles was also proposed as other mechanisms for spindle termination controlling the spindle duration. Similar to previous studies in our model ripples were generated (Brunel and Wang, 2003; Buzsaki et al., 1992; Klausberger et al., 2003; Memmesheimer, 2010; Ylinen et al., 1995) as a result of interactions between excitatory and inhibitory CA1 neurons. In spite of growing evidence for the significance of spindle-ripple coupling in memory consolidation (Binder et al., 2019; Latchoumane et al., 2017; Maingret et al., 2016; Xia et al., 2017), the mechanism underlying spindle-ripple coupling remains unclear. It has been shown that the inhibition of CA1 parvalbumin-positive cells (Xia et al., 2017) and silencing the projections from hippocampus to medial prefrontal cortex (Binder et al., 2019) disrupted the spindle-ripple coupling. Consistently in our model the long-range hippocampal-thalamocortical connections contribute to spindle-ripple coupling. Thalamic spindles can reach the hippocampus either directly by projections from the thalamus to CA1 (Vertes, 2015.) or indirectly by the prefrontal (Rajasethupathy et al., 2015) or entorhinal cortical inputs to the hippocampus (Preston and Eichenbaum, 2013). The excitation provided by the depolarizing phase of the spindle can generate the sharp wave ripples in the hippocampal network phase-locked to the spindles. We further showed that the larger temporal depolarizing window provided by the slower spindles resulted in stronger spindle-ripple coupling. On the other hand, the depolarizing inputs that the thalamocortical network receives from the hippocampus during sharp wave ripples co-occurring with spindles can increase the spindle-ripple coupling by changing the phase of the spindle towards more depolarizing phases.

To conclude, using simultaneous thalamic and hippocampal recordings from naturally sleeping mice before and after exploration, we provide the first evidence for the strong coupling of CA1 units to thalamic spindles and increase of this coupling following spatial experience. Our experimental and computational findings suggest that low-frequency long-duration thalamic spindles enhance multi-regional temporal coordination by providing large enough window of opportunity for information processing during sleep. These results shed light on our understanding of the mechanisms underlying the hippocampal-thalamocortical dialogue required for memory consolidation.

## Materials and methods

### Mice

We used a publically available dataset (Peyrache et al., 2015a, 2015b) recorded in Professor Gyorgy Buzsaki’s lab and taken from the public data sharing repository http://crcns.org. This dataset consists of 18 recording sessions (four male mice weighing ∼30 g, 3–6 months) each with 6 recording sites in CA1 and 64 recording site (8 shanks, separated by 200 μm) in anterior thalamus. Each session consists of successive epochs of baseline sleep, spatial exploration (food foraging in a circular arena) for 30 min, and post exploration-sleep.

### Electrophysiological Recordings

Electrophysiological signals were acquired at 20 kHz on a 256-channel Amplipex system (16-bit resolution) and were down sampled to 1.25 kHz for LFP analyses. SWS was detected using CA1 LFP spectrogram (Viejo and Peyrache, 2020) as periods with high delta and spindle activity during sleep (a long period of immobility tracked with the LEDs). All the analyses were conducted during SWS.

### Detection of Sleep Spindles, Ripples, and Slow Oscillations

The CA1 and thalamic LFPs were used to detect ripples and SO/spindles respectively. The algorithm for detecting ripples was similar to a previous paper (Levenstein et al., 2019). The CA1 LFP was first band pass filtered in the ripple frequency range (150–200 Hz, 4th order zero-phase delay Butterworth). The RMS signal was calculated and smoothed using a Gaussian window (50 ms). A ripple event was identified when the smoothed RMS signal was higher than median + 4 SD for a minimum duration of 30 ms. The time of the ripple event was defined as the time of the largest ripple peak. The algorithm for detecting spindles was similar to the previous study (Klinzing et al., 2016). Thalamic LFP signal was first band pass filtered in the spindle frequency range (7-10 Hz for slow spindle and 11-15 Hz for fast spindle) using finite-impulse-response (FIR) filters from the EEGLAB toolbox (Delorme and Makeig, 2004, FIR band-pass filter, filter order corresponds to 3 cycles of the low frequency cut off). Hilbert transform was used to compute instantaneous amplitude which was then smoothened using a 300 ms Gaussian window. Periods with amplitude larger than 3 SD for a duration greater than 0.5 s and lower than 3 s were selected as spindle events. Two nearby events were merged if they were closer than 0.5 s. The time of the spindle event was defined as the time of the largest spindle peak. Since in this study we sought to investigate the differential properties of slow versus fast spindles, we considered only isolated fast and slow spindles excluding the fast (slow) spindles that were accompanied by slow (fast) spindles (search window of ± 0.5 s around the spindle onset/offset).

The FMA toolbox (Zugaro et al., 2018) was used to detect SOs. First, the signal was filtered between 0.5 and 4 Hz (FIR band-pass filter, filter order corresponds to 3 cycles of the low frequency cut off). SOs were identified in the LFP when (i) the distance between consecutive positive-to-negative zero crossings was between 0.5 and 2 s (corresponding to 0.5 to 2 Hz), (ii) positive peaks greater than 2 SD were identified as SO positive peaks, and (iii) difference between the positive and negative peaks was greater than 3.5 SD.

### Phase Amplitude Coupling

The method to calculate phase amplitude coupling was similar to the previous papers (Staresina et al., 2015; Dehnavi et al., 2021). First time–frequency representations (TFRs) were calculated for every spindle event by mtmconvol function of the FieldTrip toolbox (Oostenveld et al., 2011) with frequency steps of 0.25 Hz in the frequency range of 5-300 Hz. Sliding (10 ms steps) Hanning tapered window with a variable length including five cycles were used to ensure reliable power estimates. TFRs of all epochs around an event were then normalized as percentage change from the pre-event baseline (−2.5 to −1.5 s). To quantify the modulation of hippocampal ripple amplitude with the phase of thalamic spindle, first TFR bins around each spindle were averaged across the ripple frequency ranges to obtain the power of ripple time series (−3 to 3 s around the spindle peak). To ensure proper phase estimation, both LFP and power of the ripple time series were filtered in the spindle frequency range (slow and fast spindles: 7-10 Hz and 11–15 Hz respectively, two-pass FIR band pass filter, order = 3 cycles of the low frequency cut-off). Next, we extracted the phase values of these time series using the Hilbert transform and defined a synchronization index (SI) for each spindle event as the vector mean of phase difference between the two time series.

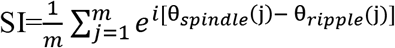

where m is the number of time points, θ_*spindle*_(j) represents the phase value of the spindle time series at time point j and θ_*ripple*_(j) shows the phase value of the fluctuations in the ripple power time series at time point j. The used interval for estimating the preferred phase was −0.25 s to +0.25 s around the spindle peak. The resulting SI is a complex number with the angle representing the phase shift between the two oscillatory events (i.e. ripple power and spindle) while the absolute value of SI represents the strength of the coupling between the two oscillatory events.

### Phase-locking analyses

To find the phase of the fast and slow spindles at the time of the unit spikes/ripples, first the instantaneous phase of thalamic LFP filtered in the fast and slow spindle frequency ranges was estimated by Hilbert transform. Then the circular mean of the phases of the spikes occurring ± 0.5 s around the spindle peaks and phases of the ripples occurring between the spindle onset and offset were computed. To obtain the phase-locking of the ripples/spindles to SO, the circular mean of the instantaneous phases of thalamic LFP filtered in the SO frequency range at the time of the ripple/spindle was computed.

### Modulation index

To examine neural spiking activity around spindles/ripples, perievent spike histograms (PETHs) were generated ±2 s around the each event and smoothed with a Gaussian window of 20 ms. Similar to a previous paper (Yang et al., 2019), the PETHs around the true events were z-score normalized to the PETHs around surrogate events obtained by randomly distributing (repeated 100 times) the same number of events detected every 4 s in a given session. The time series of surrogate PETHs were subtracted from the corresponding values of the true PETHs and the resulting PETHs were then z-score normalized to the firing rate during the entire 4 s time window of the surrogate PETHs. To quantify modulation of neural spiking to the spindle/ripple events, a modulation index (MI) was defined as the difference between peak to trough firing rate in a 200 ms window centered at 0 on the PETHs.

### Event correlation histogram

To analyze the temporal relationships between spindles and ripples, event (ripple) correlation histograms were calculated within ±1.5 s window around the reference event (spindle onset) with a bin size of 100 ms. The event correlation histograms were then normalized to a 1-s pre-event interval (form −2.0 s to −1.0 s prior to the reference event at 0 s). For statistical analysis a randomization procedure similar to a previous paper (Mölle et al., 2011) was applied by randomizing the time points of all detected ripples (repeated 100 times). Next, for each session, the event correlation histograms for the randomized data were recalculated and finally each bin in the true event correlation histogram were compared with the corresponding bin in the random condition using two-tailed paired-samples t test.

### Hippocampal-thalamocortical model

The neural mass hippocampal-thalamocortical model consists of two hippocampal networks representing CA1 and CA3 networks, one thalamic and two cortical networks. The networks were described in our previous papers (Ghorbani et al., 2012; Hashemi et al., 2019; Azimi et al., 2021). In short, each of cortical and hippocampal networks consists of one group of identical inhibitory (I) neurons (number of neurons: 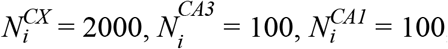, CX represents the cortical network) and one group of identical excitatory (E) neurons (number of neurons: 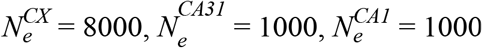). The model is a firing rate model (Ghorbani et al., 2012; Hashemi et al., 2019) so that the firing rate (*r*) of hippocampal and cortical neurons is simply a sigmoid function of the membrane potential 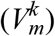.

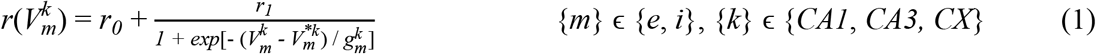

Here, *r*_1_ = 70 Hz (*r*_0_ = 0.1 Hz) is the maximal (minimum) firing rate, 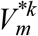 is the threshold firing potential and 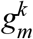 shows the sharpness of the firing rate dependence on the membrane potential (Supplementary file 1 Table 1).

The thalamic network consists of one group of identical excitatory neurons representing the thalamocortical neurons (TC) and one group of identical inhibitory neurons representing the thalamic reticular neurons (RE, number of neurons: 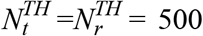, TH shows the thalamic network, t and r show TC and RE neurons respectively). A bursting variable, *u*_*m*_ is defined for thalamic neurons to model T-type calcium currents so that they can show both tonic and burst modes. As was described in our previous paper (Hashemi et al., 2019), the rate equations for thalamic neurons are given by:

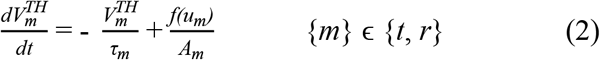

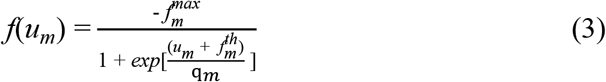

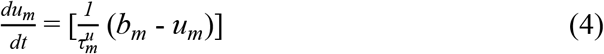

If 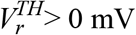 then *b*_*r*_ = 0 otherwise *b*_*r*_ = −200 mA

If 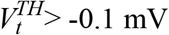 then *b*_*t*_ = 0 otherwise *b*_*t*_ = −200 mA

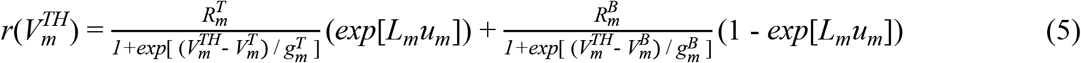

where *A*_*m*_ is the specific membrane capacitance, 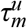 (0.22 s and 0.11 s for TC and RE neurons respectively) is the time constant that controls the dynamics of *u*_*m*_ which is much larger than the membrane time constant *τ*_*m*_ = 10 ms and *b*_*m*_ is the equilibrium value of *u*_*m*_ which has different values when the neuron is depolarized or hyperpolarized. 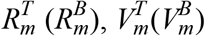 and 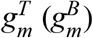 show the maximum firing rate, threshold firing potential and the sharpness of the firing rate dependence on the membrane potential during tonic (burst) mode respectively (Supplementary file 1 Table 1). The within and between network connections are considered based on anatomical connections (Supplementary file 1 Table 2). The probability and strength of connections from neuron m of network k to the neuron n of network h are given by 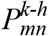 and 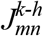 respectively. *k* and *h* can take CA1, CA3, CX or TH. For short-range connections for which h and k are the same only one of them is shown. *n* and m can be t, r, e (representing cortical or hippocampal excitatory neurons) and i (representing cortical or hippocampal inhibitory neurons). Dendritic spike frequency adaptation, (Ghorbani et al., 2012) is considered for cortical and hippocampal excitatory to excitatory connection so that the strength of these connections is a decreasing sigmoid function of an adaptation variable, c.

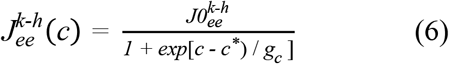

where 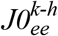 and *c*^*^ = 10 are the maximal synaptic strength and the threshold adaptation level respectively. *g*_*c*_ = 3 shows the sharpness of the excitatory to excitatory connection dependence on the adaptation variable. The parameters of the second cortical network are shown by primes.

The full model consists of below rate equations:

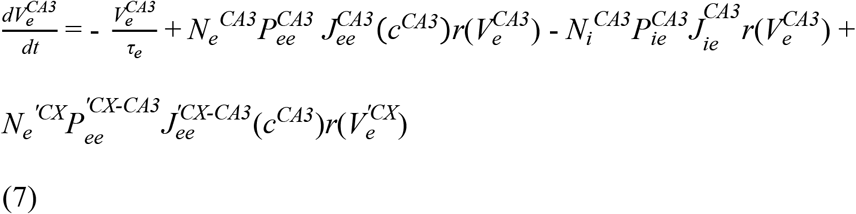

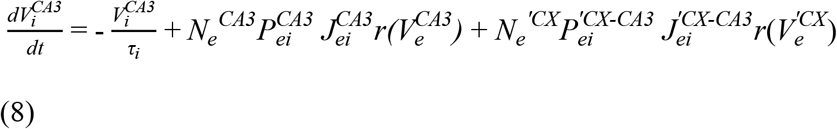

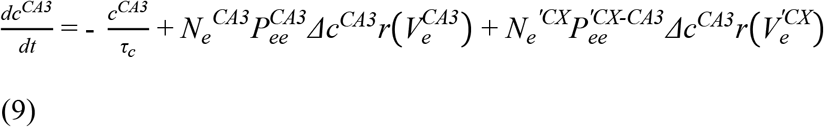

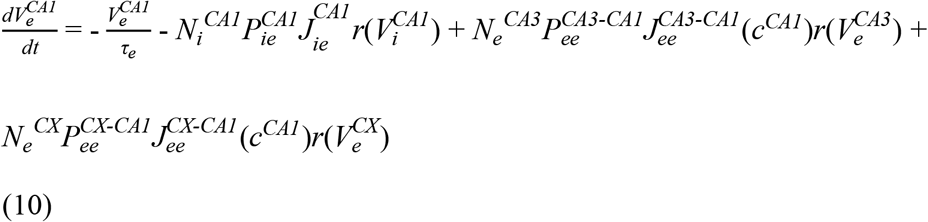

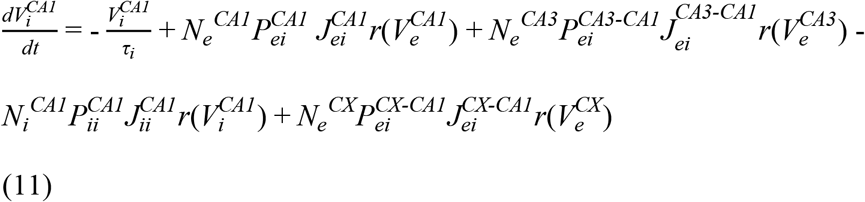

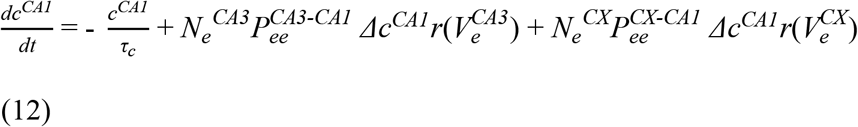

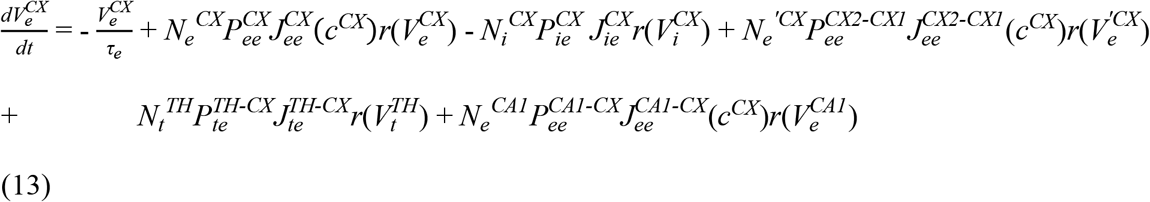

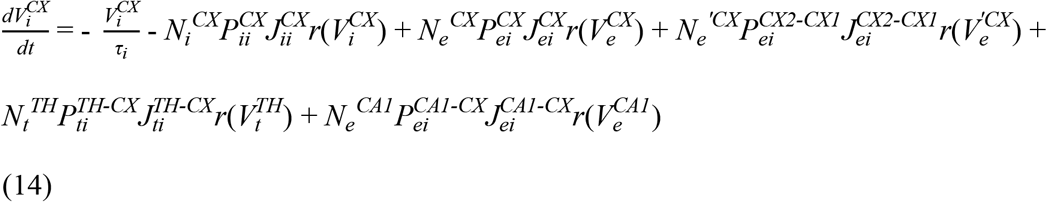

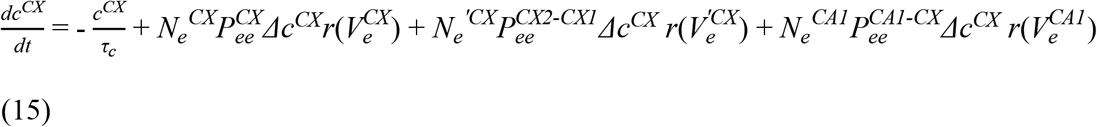

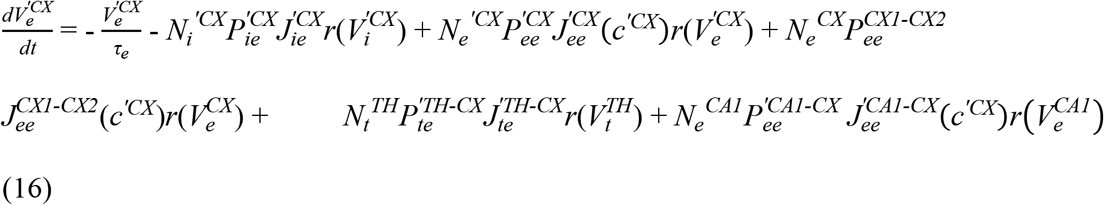

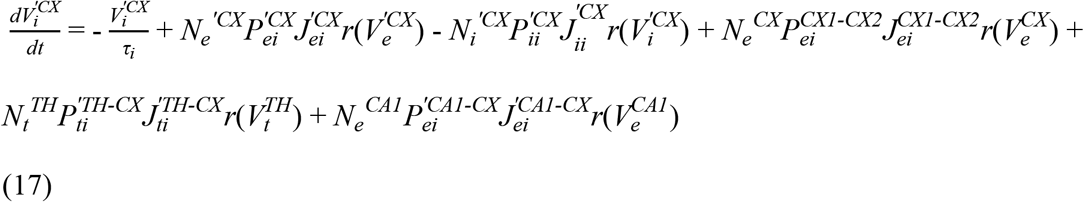

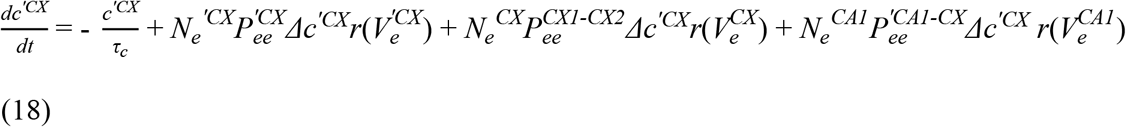

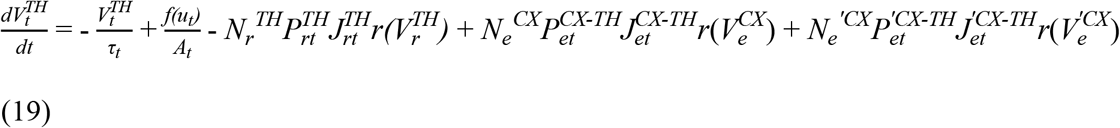

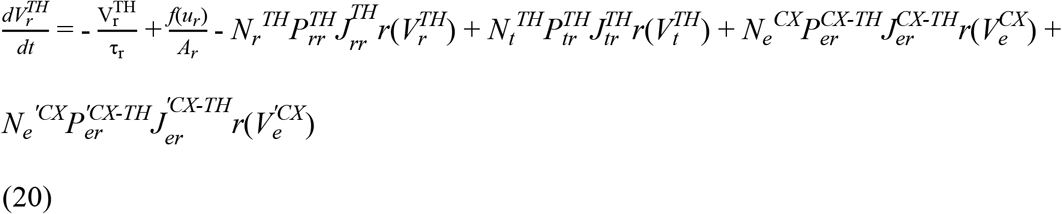

Here *τ*_*e*_ = 20 ms and *τ*_*i*_ = 10 ms are membrane time constants of excitatory and inhibitory neurons respectively, *τ*_*c*_ = 500 ms is the adaptation time constant, *Δc*^*n*^ shows the increase in the adaptation variable of neuron n as a result of excitatory to excitatory synaptic input (Supplementary file 1 Table 2).

The nonlinear equations of the model were solved using the fourth-order Runge–Kutta method with a time step of 1 ms by MATLAB R2017b software.

### Statistical Analysis

Unless otherwise stated MATLAB (version 2017b) was used for all statistical analyses. Particularly, MATLAB CircStat toolbox (Berens, 2009) was used for all statistical analyses on circular data. We applied the basic Rayleigh test (Batschelet, 1981) to investigate circular non-uniformity of the phases. circular-mtest was used for comparison of mean phases of paired circular data. Circular-linear correlation was used to compute the correlations between SO-spindle phase and spindle-ripple coupling strength. Non-circular data were compared using non-parametric tests. Mann–Whitney test and Kruskall-Wallis test (using SPSS version 26) with post hoc Dunn’s multiple correction were used for unpaired data (indicated in the text for each analysis). Wilcoxon signed rank test and Friedman test (using SPSS version 26) followed by post hoc Wilcoxon signed rank test were used for paired data.

## Data and code availability

The used experimental data sets were taken from the public CRCNS Data Sharing, doi: 10.6080/K0G15XS1. The code which is used to generate the model data is uploaded as a Source Code File (Figure 7—source code 1).

## Acknowledgments

We would like to thank Prof. Adrien Peyrache and Fereshteh Dehnavi for helpful discussions. We also thank Prof. Adrien Peyrache and Prof. Gyorgy Buzsaki for sharing their data sets.

## Author contributions

Conceptualization, M.G.; Methodology, Z.A., A.A., and M.G.; Formal analysis, Z.A., and A.A.; Writing—Original draft, Z.A., A.A., and M.G.; Writing—Review & Editing, M.G., and Z.A.; Supervision, M.G.

## Additional files

### Source code 1

Matlab code for generating the model data

### Supplementary file 1

Supplemental tables

**Transparent reporting form**

---

**Figure 4-Figure supplement 1.**
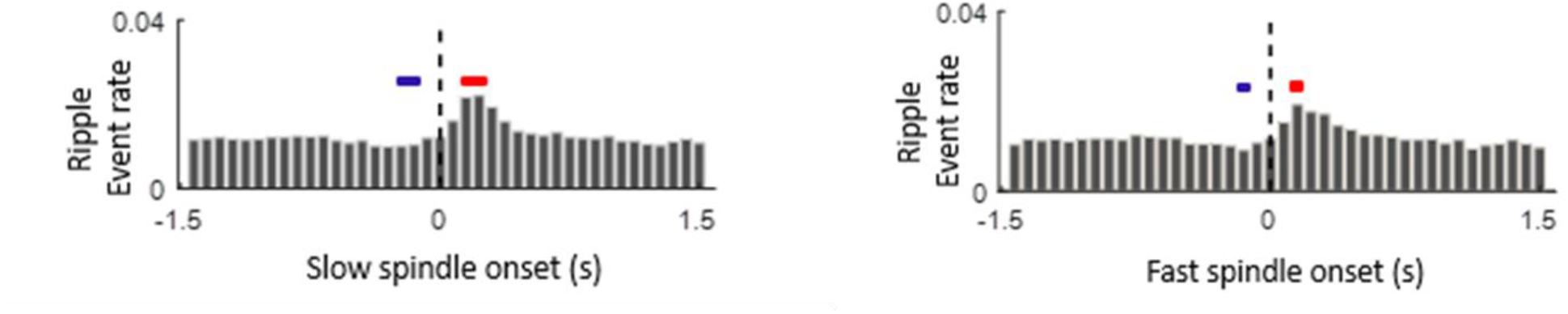
The increase of ripple occurrence after the spindle onset. The event correlation histograms of ripple events time-locked to the spindle onset for slow (Left) and fast (Right) spindles. Thick lines show the significant (p<0.05) increase (red) or decrease (blue) in ripple rates (pairwise comparison with the randomized data; see Materials and methods for details).

**Figure 5-Figure supplement 1.**
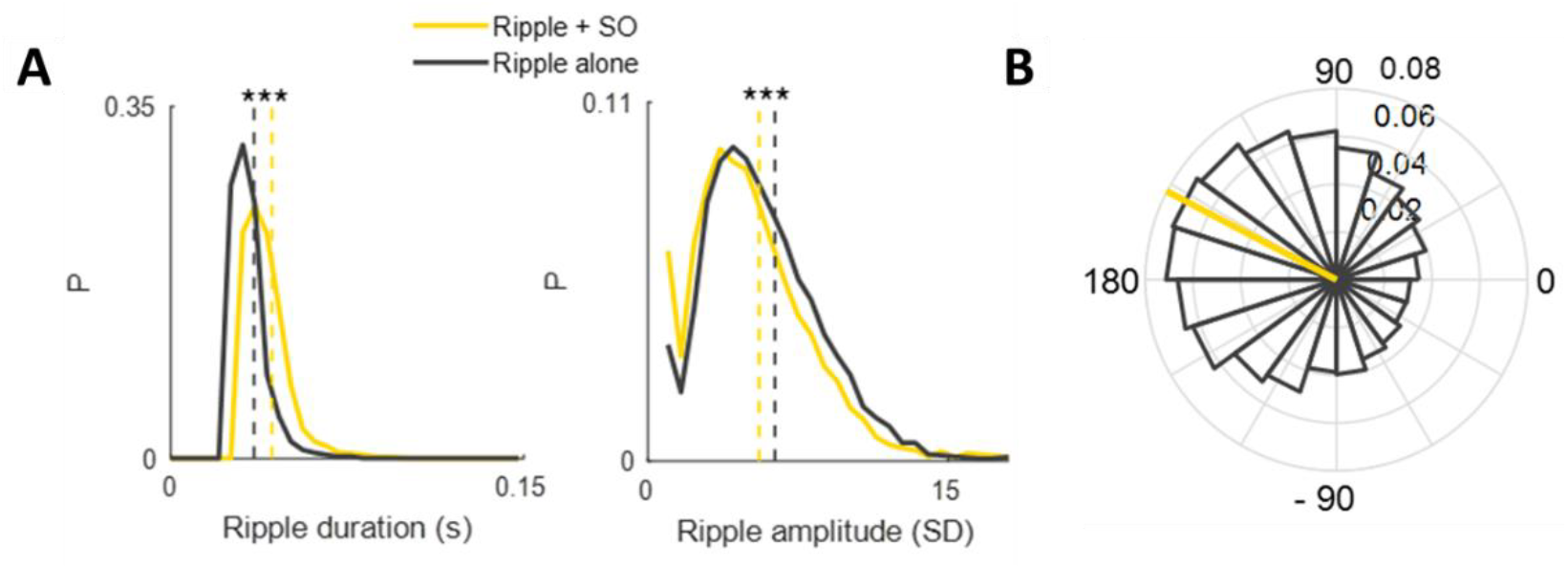
Temporal associations between SOs and ripples. **(A)** The histogram of the duration (Left) and amplitude (Right) of the ripples co-occurring with SOs (yellow, SO peak occurred within ± 0.5 s interval around ripple maximum peak) and isolated ripples (black). The vertical dotted lines show the mean values. *** p<0.001 **(B)** The circular histogram of the SO phases at the time of the ripples revealing a non-uniform distribution (p <10^−100^, z = 674.04, Rayleigh test).

**Figure 5-Figure supplement 2.**
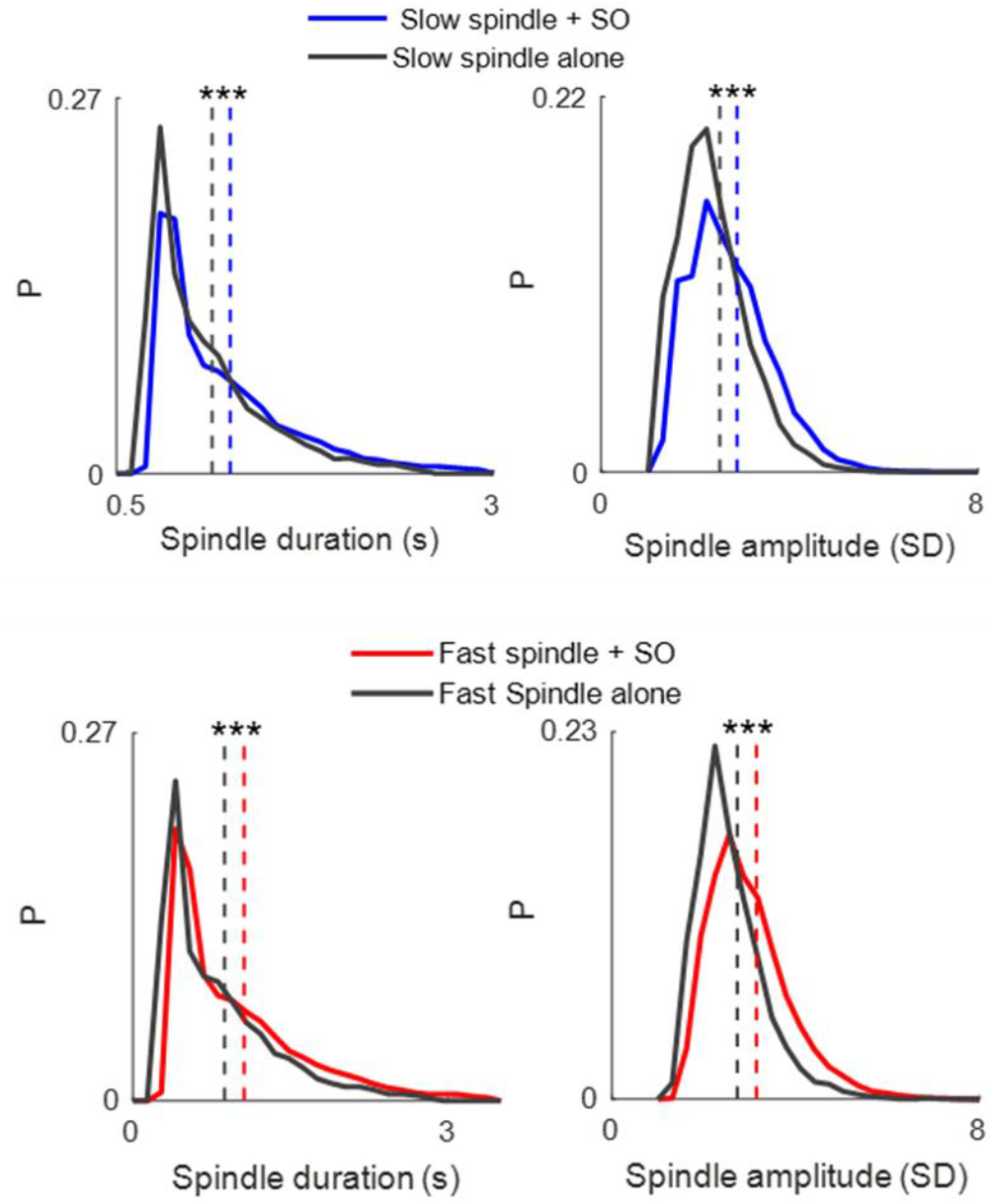
Differential properties of spindles co-occurring with SOs compared to the isolated spindles. The histogram of the duration (Left) and amplitude (Right) of the spindles co-occurring with SOs (SO peak occurred within a ± 1.5 s interval around spindle maximum peak) and isolated spindles (black) for slow (Top) and fast (Bottom) spindles. The vertical dotted lines show the mean values. *** p<0.001.

**Figure 7-Figure supplement 1.**
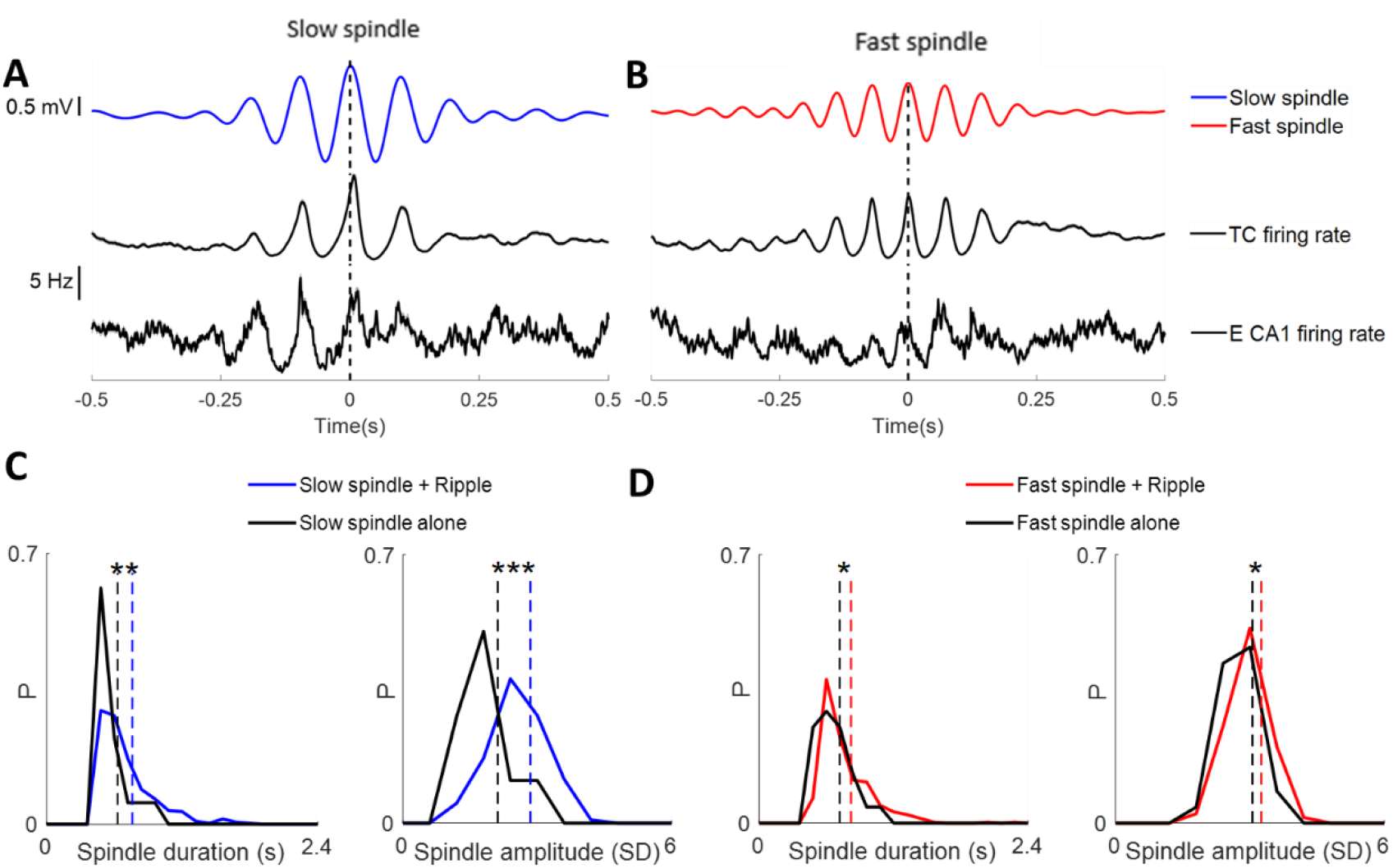
Hippocampal-thalamocortical temporal interactions during spindles in the model data. **(A)** Average membrane potential of the thalamocortical neurons filtered in the slow spindle frequency range (8-12 Hz) aligned to the spindle peaks (Top). Average of slow spindle peak– locked firing rate of thalamocortical (Middle) and CA1 excitatory (Bottom) neurons. **(B)** The same as (A) for fast spindles (12-16 Hz). **(C)** Histogram of the duration (Left) and amplitude (Right) of slow spindles co-occurring with ripples (blue) and isolated spindles. **(D)** The same as (C) for fast spindles (red).

## References

Andrillon T, Nir Y, Staba RJ, Ferrarelli F, Cirelli C, Tononi G, et al. Sleep spindles in humans: insights from intracranial EEG and unit recordings. J Neurosci. 2011;31(49):17821–34. doi:10.1523/jneurosci.2604-11.2011

Ayoub A, Aumann D, Hörschelmann A, Kouchekmanesch A, Paul P, Born J, et al. Differential effects on fast and slow spindle activity, and the sleep slow oscillation in humans with carbamazepine and flunarizine to antagonize voltage-dependent Na+ and Ca2+ channel activity. Sleep. 2013;36(6):905–11. doi:10.5665/sleep.2722

Azimi A, Alizadeh Z, Ghorbani M. The essential role of hippocampo-cortical connections in temporal coordination of spindles and ripples. NeuroImage. 2021:118485. doi:10.1016/j.neuroimage.2021.118485

Bal T, McCormick DA. What stops synchronized thalamocortical oscillations? Neuron. 1996;17(2):297–308. doi:10.1016/s0896-6273(00)80161-0

Bal T, von Krosigk M, McCormick DA. Synaptic and membrane mechanisms underlying synchronized oscillations in the ferret lateral geniculate nucleus in vitro. The Journal of physiology. 1995;483 ( Pt 3):(Pt 3):641–63. doi:10.1113/jphysiol.1995.sp020612

Bandarabadi M, Herrera CG, Gent TC, Bassetti C, Schindler K, Adamantidis AR. A role for spindles in the onset of rapid eye movement sleep. Nat Commun. 2020;11(1):5247. doi:10.1038/s41467-020-19076-2

Barthó P, Slézia A, Mátyás F, Faradzs-Zade L, Ulbert I, Harris KD, et al. Ongoing network state controls the length of sleep spindles via inhibitory activity. Neuron. 2014;82(6):1367–79. doi:10.1016/j.neuron.2014.04.046

Batschelet E. Circular Statistics in Biology. Academic Press. 1981.

Berens P. CircStat: a MATLAB toolbox for circular statistics. Journal of statistical software 2009;31(1):1–21. doi.:10.18637/jss.v031.i10.

Binder S, Mölle M, Lippert M, Bruder R, Aksamaz S, Ohl F, et al. Monosynaptic Hippocampal-Prefrontal Projections Contribute to Spatial Memory Consolidation in Mice. J Neurosci. 2019;39(35):6978–91. doi:10.1523/jneurosci.2158-18.2019

Bonjean M, Baker T, Lemieux M, Timofeev I, Sejnowski T, Bazhenov M. Corticothalamic feedback controls sleep spindle duration in vivo. J Neurosci. 2011;31(25):9124–34. doi:10.1523/jneurosci.0077-11.2011

Brunel N, Wang XJ. What determines the frequency of fast network oscillations with irregular neural discharges? I. Synaptic dynamics and excitation-inhibition balance. J Neurophysiol. 2003;90(1):415–30. doi:10.1152/jn.01095.2002

Buzsáki G, Horváth Z, Urioste R, Hetke J, Wise K. High-frequency network oscillation in the hippocampus. Science. 1992;256(5059):1025–7. doi:10.1126/science.1589772

Chatburn A, Coussens S, Lushington K, Kennedy D, Baumert M, Kohler M. Sleep spindle activity and cognitive performance in healthy children. Sleep. 2013;36(2):237–43. doi:10.5665/sleep.2380

Clemens Z, Mölle M, Eross L, Jakus R, Rásonyi G, Halász P, et al. Fine-tuned coupling between human parahippocampal ripples and sleep spindles. Eur J Neurosci. 2011;33(3):511–20. doi:10.1111/j.1460-9568.2010.07505.x

Cona F, Lacanna M, Ursino M. A thalamo-cortical neural mass model for the simulation of brain rhythms during sleep. Journal of computational neuroscience. 2014;37(1):125–48. doi:10.1007/s10827-013-0493-1

Dehnavi F, Koo-Poeggel PC, Ghorbani M, Marshall L. Spontaneous slow oscillation - slow spindle features predict induced overnight memory retention. Sleep. 2021. doi:10.1093/sleep/zsab127

Delorme A, Makeig S. EEGLAB: an open source toolbox for analysis of single-trial EEG dynamics including independent component analysis. J Neurosci Methods. 2004;134(1):9–21. doi:10.1016/j.jneumeth.2003.10.009

Destexhe A, Bal T, McCormick DA, Sejnowski TJ. Ionic mechanisms underlying synchronized oscillations and propagating waves in a model of ferret thalamic slices. J Neurophysiol. 1996;76(3):2049–70. doi:10.1152/jn.1996.76.3.2049

Destexhe A, Sejnowski TJ. Interactions between membrane conductances underlying thalamocortical slow-wave oscillations. Physiological reviews. 2003;83(4):1401–53. doi:10.1152/physrev.00012.2003

Diekelmann S, Born J. The memory function of sleep. Nature reviews Neuroscience. 2010;11(2):114–26. doi:10.1038/nrn2762

Fernandez LMJ, Lüthi A. Sleep Spindles: Mechanisms and Functions. Physiological reviews. 2020;100(2):805–68. doi:10.1152/physrev.00042.2018

Fernández-Ruiz A, Oliva A, Fermino de Oliveira E, Rocha-Almeida F, Tingley D, Buzsáki G. Long-duration hippocampal sharp wave ripples improve memory. Science. 2019;364(6445):1082–6. doi:10.1126/science.aax0758

Ferrarelli F, Huber R, Peterson MJ, Massimini M, Murphy M, Riedner BA, et al. Reduced sleep spindle activity in schizophrenia patients. Am J Psychiatry. 2007;164(3):483–92. doi:10.1176/ajp.2007.164.3.483

Fogel SM, Smith CT. The function of the sleep spindle: a physiological index of intelligence and a mechanism for sleep-dependent memory consolidation. Neurosci Biobehav Rev. 2011;35(5):1154–65. doi:10.1016/j.neubiorev.2010.12.003

Fogel SM, Smith CT, Cote KA. Dissociable learning-dependent changes in REM and non-REM sleep in declarative and procedural memory systems. Behavioural brain research. 2007;180(1):48–61. doi:10.1016/j.bbr.2007.02.037

Gardner RJ, Hughes SW, Jones MW. Differential spike timing and phase dynamics of reticular thalamic and prefrontal cortical neuronal populations during sleep spindles. J Neurosci. 2013;33(47):18469–80. doi:10.1523/jneurosci.2197-13.2013

Geiger A, Huber R, Kurth S, Ringli M, Jenni OG, Achermann P. The sleep EEG as a marker of intellectual ability in school age children. Sleep. 2011;34(2):181–9. doi:10.1093/sleep/34.2.181

Ghorbani M, Mehta M, Bruinsma R, Levine AJ. Nonlinear-dynamics theory of up-down transitions in neocortical neural networks. Physical Review E. 2012;85(2):021908. doi.org/10.1103/PhysRevE.85.021908

Hashemi NS, Dehnavi F, Moghimi S, Ghorbani M. Slow spindles are associated with cortical high frequency activity. Neuroimage. 2019;189:71–84. doi:10.1016/j.neuroimage.2019.01.012

Helfrich RF, Lendner JD, Mander BA, Guillen H, Paff M, Mnatsakanyan L, et al. Bidirectional prefrontal-hippocampal dynamics organize information transfer during sleep in humans. Nat Commun. 2019;10(1):3572. doi:10.1038/s41467-019-11444-x

Helfrich RF, Mander BA, Jagust WJ, Knight RT, Walker MP. Old Brains Come Uncoupled in Sleep: Slow Wave-Spindle Synchrony, Brain Atrophy, and Forgetting. Neuron. 2018;97(1):221–30.e4. doi:10.1016/j.neuron.2017.11.020

Jiang X, Gonzalez-Martinez J, Halgren E. Posterior Hippocampal Spindle Ripples Co-occur with Neocortical Theta Bursts and Downstates-Upstates, and Phase-Lock with Parietal Spindles during NREM Sleep in Humans. J Neurosci. 2019;39(45):8949–68. doi:10.1523/jneurosci.2858-18.2019

Kim D, Hwang E, Lee M, Sung H, Choi JH. Characterization of topographically specific sleep spindles in mice. Sleep. 2015;38(1):85–96. doi:10.5665/sleep.4330

Kim GB, Rincon Fernandez Pacheco D, Saxon D, Yang A, Sabet S, Dutra-Clarke M, et al. Rapid Generation of Somatic Mouse Mosaics with Locus-Specific, Stably Integrated Transgenic Elements. Cell. 2019;179(1):251–67.e24. doi:10.1016/j.cell.2019.08.013

Kim U, McCormick DA. The functional influence of burst and tonic firing mode on synaptic interactions in the thalamus. J Neurosci. 1998;18(22):9500–16. doi:10.1523/jneurosci.18-22-09500.1998

Klausberger T, Magill PJ, Márton LF, Roberts JD, Cobden PM, Buzsáki G, et al. Brain-state- and cell-type-specific firing of hippocampal interneurons in vivo. Nature. 2003;421(6925):844–8. doi:10.1038/nature01374

Klinzing JG, Mölle M, Weber F, Supp G, Hipp JF, Engel AK, et al. Spindle activity phase-locked to sleep slow oscillations. Neuroimage. 2016;134:607–16. doi:10.1016/j.neuroimage.2016.04.031

Ladenbauer J, Ladenbauer J, Külzow N, de Boor R, Avramova E, Grittner U, et al. Promoting Sleep Oscillations and Their Functional Coupling by Transcranial Stimulation Enhances Memory Consolidation in Mild Cognitive Impairment. J Neurosci. 2017;37(30):7111–24. doi:10.1523/jneurosci.0260-17.2017

Latchoumane CV, Ngo HV, Born J, Shin HS. Thalamic Spindles Promote Memory Formation during Sleep through Triple Phase-Locking of Cortical, Thalamic, and Hippocampal Rhythms. Neuron. 2017;95(2):424–35.e6. doi:10.1016/j.neuron.2017.06.025

Levenstein D, Buzsáki G, Rinzel J. NREM sleep in the rodent neocortex and hippocampus reflects excitable dynamics. Nat Commun. 2019;10(1):2478. doi:10.1038/s41467-019-10327-5

Lüthi A, McCormick DA. Periodicity of thalamic synchronized oscillations: the role of Ca2+− mediated upregulation of Ih. Neuron. 1998;20(3):553–63. doi:10.1016/s0896-6273(00)80994-0

Lüthi A, Bal T, McCormick DA. Periodicity of thalamic spindle waves is abolished by ZD7288,a blocker of Ih. J Neurophysiol. 1998;79(6):3284–9. doi:10.1152/jn.1998.79.6.3284

Maingret N, Girardeau G, Todorova R, Goutierre M, Zugaro M. Hippocampo-cortical coupling mediates memory consolidation during sleep. Nat Neurosci. 2016;19(7):959–64. doi:10.1038/nn.4304

Mak-McCully RA, Rolland M, Sargsyan A, Gonzalez C, Magnin M, Chauvel P, et al. Coordination of cortical and thalamic activity during non-REM sleep in humans. Nat Commun. 2017;8:15499. doi:10.1038/ncomms15499

Memmesheimer RM. Quantitative prediction of intermittent high-frequency oscillations in neural networks with supralinear dendritic interactions. Proc Natl Acad Sci U S A. 2010;107(24):11092–7. doi:10.1073/pnas.0909615107

Mölle M, Bergmann TO, Marshall L, Born J. Fast and slow spindles during the sleep slow oscillation: disparate coalescence and engagement in memory processing. Sleep. 2011;34(10):1411–21. doi:10.5665/sleep.1290

Mölle M, Eschenko O, Gais S, Sara SJ, Born J. The influence of learning on sleep slow oscillations and associated spindles and ripples in humans and rats. Eur J Neurosci. 2009;29(5):1071–81. doi:10.1111/j.1460-9568.2009.06654.x

Mölle M, Yeshenko O, Marshall L, Sara SJ, Born J. Hippocampal sharp wave-ripples linked to slow oscillations in rat slow-wave sleep. J Neurophysiol. 2006;96(1):62–70. doi:10.1152/jn.00014.2006

Morin A, Doyon J, Dostie V, Barakat M, Hadj Tahar A, Korman M, et al. Motor learning increases sleep spindles and fast frequencies in post-training sleep. Sleep. Sleep. 2008;31(8):1149–56. doi:10.5665/sleep/31.8.1149

Muehlroth BE, Sander MC, Fandakova Y, Grandy TH, Rasch B, Shing YL, et al. Precise Slow Oscillation-Spindle Coupling Promotes Memory Consolidation in Younger and Older Adults. Sci Rep. 2019;9(1):1940. doi:10.1038/s41598-018-36557-z

Nicolas A, Petit D, Rompré S, Montplaisir J. Sleep spindle characteristics in healthy subjects of different age groups. Clin Neurophysiol. 2001;112(3):521–7. doi:10.1016/s1388-2457(00)00556-3

Niethard N, Ngo HV, Ehrlich I, Born J. Cortical circuit activity underlying sleep slow oscillations and spindles. Proc Natl Acad Sci U S A. 2018;115(39):E9220–e9. doi:10.1073/pnas.1805517115

Niknazar M, Krishnan GP, Bazhenov M, Mednick SC. Coupling of Thalamocortical Sleep Oscillations Are Important for Memory Consolidation in Humans. PLoS One. 2015;10(12):e0144720. doi:10.1371/journal.pone.0144720

Ngo HV, Fell J, Staresina B. Sleep spindles mediate hippocampal-neocortical coupling during long-duration ripples. Elife. 2020;9. doi:10.7554/eLife.57011

Oostenveld R, Fries P, Maris E, Schoffelen JM. FieldTrip: Open source software for advanced analysis of MEG, EEG, and invasive electrophysiological data. Comput Intell Neurosci. 2011;2011:156869. doi:10.1155/2011/156869

Oyanedel CN, Durán E, Niethard N, Inostroza M, Born J. Temporal associations between sleep slow oscillations, spindles and ripples. Eur J Neurosci. 2020;52(12):4762–78. doi:10.1111/ejn.14906

Peyrache A, Battaglia FP, Destexhe A. Inhibition recruitment in prefrontal cortex during sleep spindles and gating of hippocampal inputs. Proc Natl Acad Sci U S A. 2011;108(41):17207–12. doi:10.1073/pnas.1103612108

Peyrache, A., Lacroix, M.M., Petersen, P.C., and Buzsáki, G. (2015a). Internally organized mechanisms of the head direction sense. Nat. Neurosci. 18, 569–575. doi:10.1038/nn.3968

[dataset] Peyrache, A., Petersen P., Buzsáki, G. (2015b). Extracellular recordings from multi-site silicon probes in the anterior thalamus and subicular formation of freely moving mice. CRCNS.org. doi:10.6080/K0G15XS1

Preston AR, Eichenbaum H. Interplay of hippocampus and prefrontal cortex in memory. Curr Biol. 2013;23(17):R764–73. doi:10.1016/j.cub.2013.05.041

Rajasethupathy P, Sankaran S, Marshel JH, Kim CK, Ferenczi E, Lee SY, et al. Projections from neocortex mediate top-down control of memory retrieval. Nature. 2015;526(7575):653–9. doi:10.1038/nature15389

Siapas AG, Wilson MA. Coordinated interactions between hippocampal ripples and cortical spindles during slow-wave sleep. Neuron. 1998;21(5):1123–8. doi:10.1016/s0896-6273(00)80629-7

Sirota A, Csicsvari J, Buhl D, Buzsáki G. Communication between neocortex and hippocampus during sleep in rodents. Proc Natl Acad Sci U S A. 2003;100(4):2065–9. doi:10.1073/pnas.0437938100

Staresina BP, Bergmann TO, Bonnefond M, van der Meij R, Jensen O, Deuker L, et al. Hierarchical nesting of slow oscillations, spindles and ripples in the human hippocampus during sleep. Nat Neurosci. 2015;18(11):1679–86. doi:10.1038/nn.4119

Sullivan D, Mizuseki K, Sorgi A, Buzsáki G. Comparison of sleep spindles and theta oscillations in the hippocampus. J Neurosci. 2014;34(2):662–74. doi:10.1523/jneurosci.0552-13.2014

Taillard J, Sagaspe P, Berthomier C, Brandewinder M, Amieva H, Dartigues JF, et al. Non-REM Sleep Characteristics Predict Early Cognitive Impairment in an Aging Population. Frontiers in neurology. 2019;10:197. doi:10.3389/fneur.2019.00197

Timofeev, I., Bazhenov, M. Mechanisms and biological role of thalamocortical oscillations. Trends in Chronobiology Research. 2005:1–47.

Varela C, Wilson MA. mPFC spindle cycles organize sparse thalamic activation and recently active CA1 cells during non-REM sleep. Elife. 2020;9. doi:10.7554/eLife.48881

Vertes RP. Major diencephalic inputs to the hippocampus: supramammillary nucleus and nucleus reuniens. Circuitry and function. Progress in brain research. 2015;219:121–44. doi:10.1016/bs.pbr.2015.03.008

Viejo G, Peyrache A. Precise coupling of the thalamic head-direction system to hippocampal ripples. Nat Commun. 2020;11(1):2524. doi:10.1038/s41467-020-15842-4

Wierzynski CM, Lubenov EV, Gu M, Siapas AG. State-dependent spike-timing relationships between hippocampal and prefrontal circuits during sleep. Neuron. 2009;61(4):587–96. doi:10.1016/j.neuron.2009.01.011

Xia F, Richards BA, Tran MM, Josselyn SA, Takehara-Nishiuchi K, Frankland PW. Parvalbumin-positive interneurons mediate neocortical-hippocampal interactions that are necessary for memory consolidation. Elife. 2017;6. doi:10.7554/eLife.27868

Yuan RK, Lopez MR, Ramos-Alvarez MM, Normandin ME, Thomas AS, Uygun DS, et al. Differential effect of sleep deprivation on place cell representations, sleep architecture, and memory in young and old mice. Cell reports. 2021;35(11):109234. doi:10.1016/j.celrep.2021.109234

Ylinen A, Bragin A, Nádasdy Z, Jandó G, Szabó I, Sik A, et al. Sharp wave-associated high-frequency oscillation (200 Hz) in the intact hippocampus: network and intracellular mechanisms. J Neurosci. 1995;15(1 Pt 1):30–46. doi:10.1523/jneurosci.15-01-00030.1995

Zugaro, M., Todorova, R., Girardeau, G., Cei, A., El Kanbi, K. FMAToolbox. doi:http://fmatoolbox.sourceforge.net/

